# Molecular characterization of selectively vulnerable neurons in Alzheimer’s Disease

**DOI:** 10.1101/2020.04.04.025825

**Authors:** Kun Leng, Emmy Li, Rana Eser, Antonia Piergies, Rene Sit, Michelle Tan, Norma Neff, Song Hua Li, Roberta Diehl Rodriguez, Claudia Kimie Suemoto, Renata Elaine Paraizo Leite, Carlos A. Pasqualucci, William W. Seeley, Salvatore Spina, Helmut Heinsen, Lea T. Grinberg, Martin Kampmann

**Affiliations:** Institute for Neurodegenerative Disease, University of California, San Francisco, San Francisco, CA, USA; Chan Zuckerberg Biohub, San Francisco, CA, USA; Biomedical Sciences Graduate Program, University of California, San Francisco, San Francisco, CA, USA; Medical Scientist Training Program, University of California, San Francisco, San Francisco, CA, USA; Memory and Aging Center, Weill Institute for Neurosciences, Department of Neurology, University of California, San Francisco, San Francisco, CA, USA; Department of Neurology, Universidade de São Paulo, Faculdade de Medicina, São Paulo, Brazil; Department of Pathology, Universidade de São Paulo, Faculdade de Medicina, São Paulo, Brazil; Division of Geriatrics, Department of Clinical Medicine, Universidade de São Paulo, Faculdade de Medicina, São Paulo, Brazil; Department of Psychiatry, University of Würzburg, Würzburg, Germany; Global Brain Health Institute, University of California, San Francisco, San Francisco, CA, USA; Department of Pathology, University of California, San Francisco, San Francisco, CA, USA; Department of Biochemistry and Biophysics, University of California, San Francisco, San Francisco, CA, USA

**Author notes:** These authors contributed equally: Kun Leng, Emmy Li.

## Abstract

Alzheimer’s disease (AD) is characterized by the selective vulnerability of specific neuronal populations, the molecular signatures of which are largely unknown. To identify and characterize selectively vulnerable neuronal populations, we used single-nucleus RNA sequencing to profile the caudal entorhinal cortex and the superior frontal gyrus – brain regions where neurofibrillary inclusions and neuronal loss occur early and late in AD, respectively – from postmortem brains spanning the progression of AD-type tau neurofibrillary pathology. We identified RORB as a marker of selectively vulnerable excitatory neurons in the entorhinal cortex, and subsequently validated their depletion and selective susceptibility to neurofibrillary inclusions during disease progression using quantitative neuropathological methods. We also discovered an astrocyte subpopulation, likely representing reactive astrocytes, characterized by decreased expression of genes involved in homeostatic functions. Our characterization of selectively vulnerable neurons in AD paves the way for future mechanistic studies of selective vulnerability and potential therapeutic strategies for enhancing neuronal resilience.

## MAIN TEXT

Selective vulnerability is a fundamental feature of neurodegenerative diseases, in which different neuronal populations show a gradient of susceptibility to degeneration^1, 2^. Selective vulnerability at the network level has been extensively explored in Alzheimer’s disease (AD)^3-5^, currently the leading cause of dementia and lacking in effective therapies. However, little is known about the mechanisms underlying selective vulnerability at the cellular level in AD, which could provide insight into disease mechanisms and lead to therapeutic strategies.

The entorhinal cortex (EC), an allocortex, is one of the first cortical brain regions to exhibit neuronal loss in AD^6^. Neurons in the external EC layers, especially in layer II, accumulate tau-positive neurofibrillary inclusions and die early in the course of AD^7-12^. However, these selectively vulnerable neurons have yet to be characterized extensively at the molecular level. Furthermore, it is unknown whether there are differences in vulnerability among subpopulations of these neurons. Although rodent models of AD have offered important insights^13-15^, the available models fail to capture some critical disease processes simultaneously, such as the accumulation of neurofibrillary inclusions and neuronal loss^16^, limiting the extrapolation of findings from rodent models to address selective vulnerability.

Previous studies have combined laser capture microdissection with microarray analysis of gene expression^17, 18^ to characterize EC neurons in AD, but focused on disease-related changes in gene expression, rather than selective vulnerability. More recently, single-nucleus RNA-sequencing (snRNA-seq) has enabled large-scale characterization of transcriptomic profiles of individual cells from post-mortem human brain tissue^19, 20^. However, snRNA-seq studies of AD published to date have focused on cell-type specific differential gene expression between AD cases and healthy controls^21, 22^, without explicitly addressing selective vulnerability.

Here, we performed snRNA-seq on post-mortem brain tissue from a cohort of cases spanning the progression of AD-type tau neurofibrillary pathology to characterize changes in the relative abundance of cell types and cell type subpopulations. Importantly, we discovered a selectively vulnerable subpopulation of excitatory neurons in the entorhinal cortex and validated the selective depletion of this subpopulation during AD progression with quantitative histopathology, using multiplex immunofluorescence in EC regions delineated by rigorous cytoarchitectonic criteria. In addition, we examined subpopulations of inhibitory neurons, which did not show differences in vulnerability, and also subpopulations of microglia, oligodendrocytes, and astrocytes. We uncovered an astrocyte subpopulation likely corresponding to reactive astrocytes that showed downregulation of genes involved in homeostatic function.

## RESULTS

### Cohort selection and cross-sample alignment

We performed snRNA-seq on cell nuclei extracted from postmortem brain tissue (see Methods) from the entorhinal cortex (EC) at the level of the mid-uncus and from the superior frontal gyrus (SFG) at the level of the anterior commissure (Brodmann area 8), from 10 male *APOE* ε3/ε3 individuals representing the cortical-free, early and late stages of AD-type tau neurofibrillary pathology (Braak stages^3^ 0, 2 and 6; Fig. 1a, Table 1).

**Table 1.** Description of post-mortem cohort. Asterisks denote cases used both for snRNA-seq and immunofluorescence validation. The AD neuropathological change (ADNC) score incorporates assessment of amyloid-beta deposits (“A”), staging of neurofibrillary tangles (“B”), and scoring of neuritic plaques (“C”)^100^. The Clinical Dementia Rating (CDR) reflects the degree of cognitive impairment^101^.

| Cases used for snRNA-seq |  |  |  |  |  |  |  |  |
| --- | --- | --- | --- | --- | --- | --- | --- | --- |
| Case # | Braak stage | Sex | Age at death (years) | Post-mortem interval (hours) | ADNC score | CDR before death | APOE genotype | Source |
| 1 | 0 | M | 50 | 13 | A0,B0,C0 | 0 | E3/E3 | BBAS |
| 2 | 0 | M | 60 | 12 | A0,B0,C0 | 0.5 | E3/E3 | BBAS |
| 3 | 0 | M | 71 | 12 | A1,B0,C0 | 0 | E3/E3 | BBAS |
| 4 | 2 | M | 72 | 15 | A1,B1,C0 | 0 | E3/E3 | BBAS |
| 5* | 2 | M | 77 | 4.9 | A2,B1,C1 | 0.5 | E3/E3 | UCSF |
| 6* | 2 | M | 87 | 30 | A2,B1,C2 | 2 | E3/E3 | UCSF |
| 7* | 2 | M | 91 | 50 | A1,B1,C1 | 0 | E3/E3 | UCSF |
| 8* | 6 | M | 72 | 6.9 | A3,B3,C3 | 3 | E3/E3 | UCSF |
| 9* | 6 | M | 82 | 6.7 | A3,B3,C3 | 3 | E3/E3 | UCSF |
| 10 | 6 | M | 82 | 9 | A3,B3,C3 | 3 | E3/E3 | UCSF |
| Cases used for immunofluorescence validation |  |  |  |  |  |  |  |  |
| Case # | Braak stage | Sex | Age at death | Post-mortem interval (hours) | ADNC score | CDR before death | APOE genotype | Source |
| 5* | 2 | M | 77 | 4.9 | A2,B1,C1 | 0.5 | E3/E3 | UCSF |
| 6* | 2 | M | 87 | 30 | A2,B1,C2 | 2 | E3/E3 | UCSF |
| 7* | 2 | M | 91 | 50 | A1,B1,C1 | 0 | E3/E3 | UCSF |
| 8* | 6 | M | 72 | 6.9 | A3,B3,C3 | 3 | E3/E3 | UCSF |
| 9* | 6 | M | 82 | 6.7 | A3,B3,C3 | 3 | E3/E3 | UCSF |
| 11 | 0 | F | 62 | 10.1 | A1,B0,C0 | 0 | NA | BBAS |
| 12 | 0 | M | 64 | 12 | A0,B0,C0 | 0 | E3/E3 | BBAS |
| 13 | 1 | M | 60 | 19 | A0,B1,C0 | 0 | NA | BBAS |
| 14 | 1 | F | 64 | 13 | A1,B1,C0 | 0 | E3/E3 | BBAS |
| 15 | 1 | M | 70 | 11 | A1,B1,C0 | 0 | E3/E3 | BBAS |
| 16 | 1 | F | 82 | 9.6 | A1,B1,C0 | 0 | NA | BBAS |
| 17 | 2 | F | 79 | 18 | A1,B1,C1 | 0 | E3/E3 | BBAS |
| 18 | 2 | F | 81 | 30.3 | A1,B1,C0 | NA | E3/E3 | UCSF |
| 19 | 3 | M | 81 | 8.3 | A2,B2,C3 | 1 | NA | UCSF |
| 20 | 3 | M | 84 | 28 | A3,B2,C2 | 1 | NA | UCSF |
| 21 | 3 | F | 88 | 9.8 | A3,B2,C2 | 0.5 | E3/E3 | UCSF |
| 22 | 3 | M | 89 | 9.1 | A3,B2,C2 | 1 | E3/E3 | UCSF |
| 23 | 4 | F | 87 | 9.5 | A1,B2,C3 | 2 | E3/E3 | UCSF |
| 24 | 4 | M | 91 | 11.2 | A3,B2,C2 | 0.5 | E3/E3 | UCSF |
| 25 | 4 | M | 103 | 7.8 | A1,B2,C2 | NA | E3/E3 | UCSF |
| 26 | 5 | M | 77 | 8.4 | A3,B3,C3 | 0.5 | E4/E4 | UCSF |
| 27 | 5 | M | 85 | 11.2 | A3,B3,C3 | 1 | E3/E3 | UCSF |
| 28 | 5 | M | 86 | 8.6 | A3,B3,C3 | 2 | E3/E4 | UCSF |
| 29 | 5 | F | 87 | 17 | A3,B3,C2 | 3 | E3/E3 | BBAS |
| 30 | 6 | F | 64 | 7.3 | A3,B3,C3 | 3 | E3/E4 | UCSF |
| 31 | 6 | F | 67 | 9.7 | A3,B3,C3 | 3 | E4/E4 | UCSF |

**Fig. 1.**
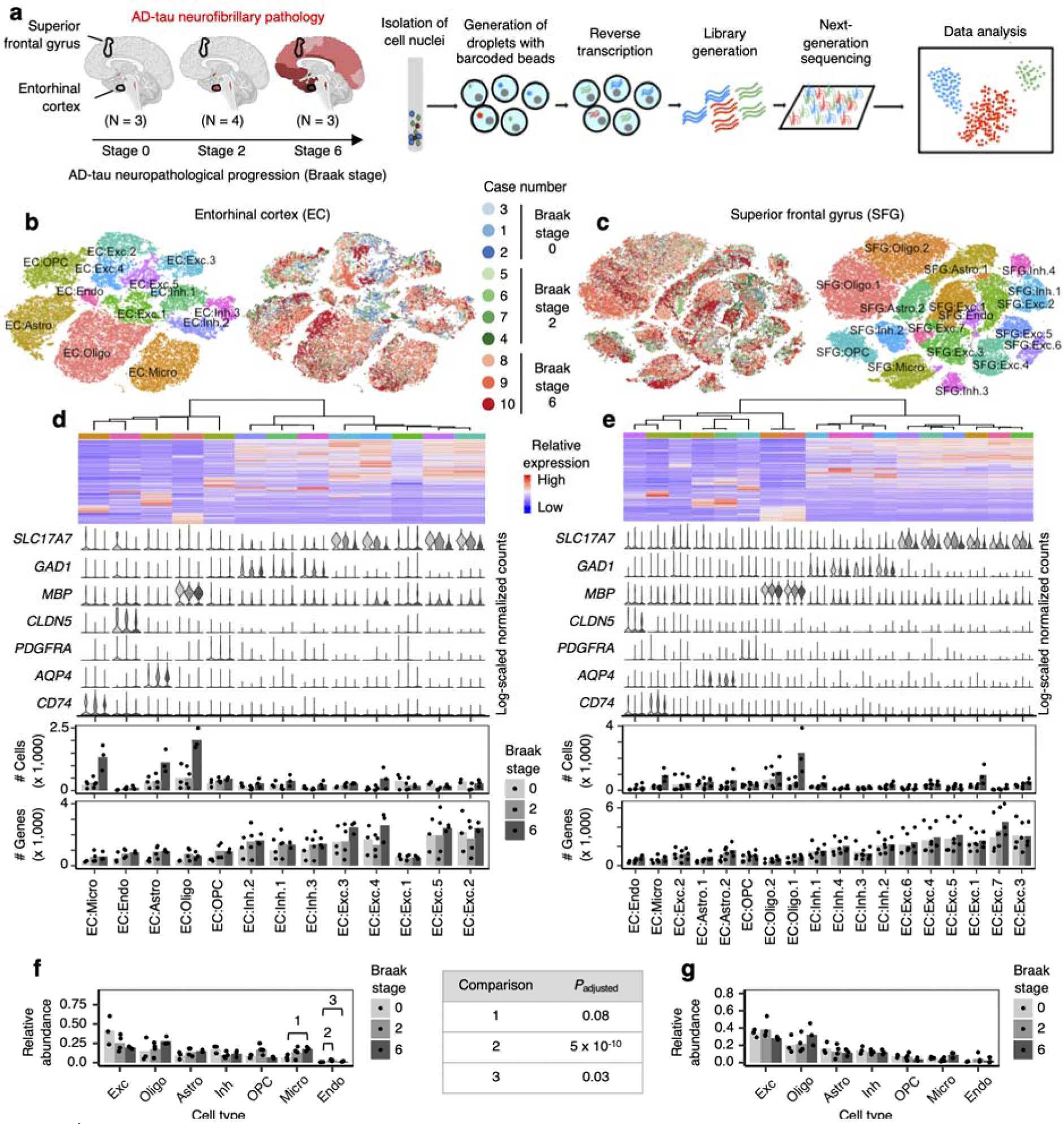
AD progression differentially affects the cell-type composition of the EC and SFG. **a**, Schematic of experimental design and sample processing. Darker shades of red in brain cartoons reflect more severe AD-tau neurofibrillary pathology. **b-c**, tSNE projection of cells from the EC (**b**) and SFG (**c**) in their respective alignment spaces, colored by individual of origin (center) or cluster assignment (outer). **d-e**, Heatmap and hierarchical clustering of clusters and cluster marker expression (top subpanel); “High” and “Low” relative expression reflect above- and below-average expression, respectively (see Methods). Expression of cell type markers in each cluster (second subpanel). The average number of cells and average number of genes detected per cell in each cluster (third and fourth subpanels). **f-g**, Relative abundance of major cell types across Braak stages. Cell type abbreviations: Exc – excitatory neurons, Oligo – oligodendrocytes, Astro – astrocytes, Inh – inhibitory neurons, OPC – oligodendrocyte precursor cells, Micro – microglia, Endo – endothelial cells.

The neuropathological hallmarks of AD are amyloid plaques, which are measured by the CERAD scores^23^ and Thal phases^24^, and neurofibrillary inclusions consisting of hyperphosphorylated tau protein (phospho-tau) aggregates, which are measured by the Braak staging system^3^. The Braak staging system is based on the stereotypical topographical progression of neurofibrillary inclusions to different brain regions. Neurofibrillary inclusions are first found in specific subcortical structures in the brainstem (Braak stages a-c, also hereon referred to collectively as Braak stage 0). Subsequently, the transentorhinal and entorhinal cortices, followed by the hippocampal formation, are the first areas of the cerebral cortex to accumulate tau pathology (Braak stages 1-2). The limbic areas and temporal neocortex then follow (Braak stages 3-4), and finally, other neocortical association areas (such as the SFG) and primary neocortical areas are involved (Braak stages 5-6)^3, 25^. Since the accumulation of neurofibrillary inclusions is the best correlate of clinical cognitive decline, after neuronal loss^26^, we reasoned that profiling matched EC and SFG samples across different Braak stages would allow us to isolate the effect of disease progression on cell types and cell type subpopulations.

A challenge in characterizing the impact of disease progression on different cell type subpopulations is that these subpopulations need to be defined in a way that is independent from the effect of disease progression. Typically, cell type subpopulations are defined by sub-grouping cells of the same cell type through cluster analysis (i.e. clustering), followed by examination of marker gene expression in the resulting clusters. To remove the effect of disease progression on clustering, we performed, prior to clustering, cross-sample alignment^27-29^ of the data from each brain region using scAlign (see Methods), which learns a low-dimensional manifold (i.e. the alignment space) in which cells tend to cluster in a manner consistent with their biological function independent of technical and experimental factors^29^. Importantly, after identifying clusters in the alignment space, we used the original data for subsequent analyses involving examination of gene expression, such as identifying differentially expressed genes between clusters.

### Changes in broad cell type composition with neuropathological AD progression

After quality control (see Methods), we recovered 42,528 cells from the EC and 63,608 cells from the SFG. Examination of the average number of genes and unique molecular identifiers (UMIs) detected per cell showed similar or superior transcript coverage compared to previously published AD snRNA-seq datasets^21, 22^ (Extended Data Fig. 1a,b).

After cross-sample alignment, we performed clustering and recovered 13 clusters in the EC and 18 clusters in the SFG. In both brain regions, clusters demonstrated spatial grouping in t-stochastic neighborhood embedding (tSNE) that was largely uncorrelated with the individual of origin (Fig. 1b,c). Furthermore, clusters showed specific expression of cell type markers and grouped in a manner consistent with their expression of cell type markers in hierarchical clustering (Fig. 1d,e, see Methods). For comparison, we also performed clustering without cross-sample alignment, which resulted in many clusters that were defined by individual of origin in addition to cell type (Extended Data Fig. 1c-f). Having confirmed the effectiveness of cross-sample alignment in removing the effect of technical and experimental factors on clustering, we then assigned clusters to broad cell types (i.e. excitatory neurons, inhibitory neurons, astrocytes, oligodendrocytes, oligodendrocyte precursor cells, microglia, and endothelial cells) based on their expression of cell type markers (Fig. 1d,e, see Methods).

Next, to assess whether the proportions of broad cell types in the EC and SFG change with disease progression, we aggregated clusters assigned to the same cell type for each individual and then computed the relative abundance of each cell type in each individual. We tested the statistical significance of changes in relative abundance using beta regression^30^ (see Methods), which is suitable for variables ranged from 0 to 1. After correcting for multiple testing (Holm’s method, threshold for significant adjusted *P*-values set at 0.05; see Methods), we found statistically significant increases in the relative abundance of endothelial cells in the EC (Fig. 1f) in Braak stages 2 and 6 compared to Braak stage 0, although the magnitude of the changes were small. We also found a trend towards increased relative abundance of microglia in the EC in Braak stage 6 (*P*_adjusted_ = 0.08), suggestive of microgliosis. In the SFG, however, we did not observe an upward trend in the relative abundance of microglia with disease progression (Fig. 1g). As for other broad cell types, we did not detect changes in relative abundance that were statistically significant after correction for multiple hypothesis testing. However, we observed a downward trend in the relative abundance of EC excitatory neurons in Braak stages 2 (*P*_unadjusted_ = 0.18) and 6 (*P*_unadjusted_ = 0.02), and of SFG excitatory neurons only in Braak stage 6 (*P*_unadjusted_ = 0.05), consistent with early involvement of the EC and sparing of the SFG until late Braak stages, and the previously described greater vulnerability of excitatory neurons relative to inhibitory neurons in AD^15, 31^.

### Selective vulnerability of excitatory neuron subpopulations

Previous single-cell transcriptomic studies of human and mouse cortex have shown that unbiased clustering of excitatory neurons largely recapitulates the laminar organization of the cortex^19, 20^. In the context of AD, tau neurofibrillary inclusions are known to preferentially accumulate in neocortical layers III and V^3, 32, 33^, most likely reflecting the selective vulnerability of specific neuronal subpopulations. Therefore, we asked whether specific excitatory neuron subpopulations show a decline in their relative abundance with disease progression, by performing subclustering of excitatory neurons in the EC and SFG after cross-sample alignment (see Methods).

In the EC, we discerned nine excitatory neuron subpopulations (Fig. 2a-d). These subpopulations exhibited distinct expression of EC layer-specific genes identified in the mouse medial EC^34^, which phylogenetically resembles the human caudal EC^35, 36^. Notably, subpopulation EC:Exc.s2 showed a striking ∼50% decrease in its relative abundance in Braak stage 2 compared to Braak stage 0, with no further decrease in Braak stage 6 (Fig. 2c), suggesting depletion early in disease. EC:Exc.s1 and EC:Exc.s4 similarly exhibited a ∼50-60% reduction in their relative abundance in Braak stage 2. EC:Exc.s1, EC:Exc.s2, and EC:Exc.s4 expressed genes associated with mouse EC layer II (Fig. 2c), consistent with the fact that tau neurofibrillary inclusions are known to accumulate preferentially in human EC layer II early in AD^7-10^. However, not all subpopulations expressing genes associated with mouse EC layer II showed similar levels of early vulnerability. For example, EC:Exc.s6 and EC:Exc.s8 did not demonstrate statistically significant changes in their relative abundance across disease progression. Outside of EC layer II, we failed to find evidence of selective vulnerability in neuronal subpopulations expressing genes associated with mouse EC layer III (EC:Exc.s0) or V/VI (EC:Exc.7, EC:Exc.s5, EC:Exc.s3). In fact, EC:Exc.s5 exhibited a statistically significant increase in its relative abundance in Braak stage 2. Since neurons are post-mitotic, this increase is likely due to the selective earlier depletion of more vulnerable excitatory neuron subpopulations, followed by later depletion of EC:Exc.s5.

**Fig. 2.**
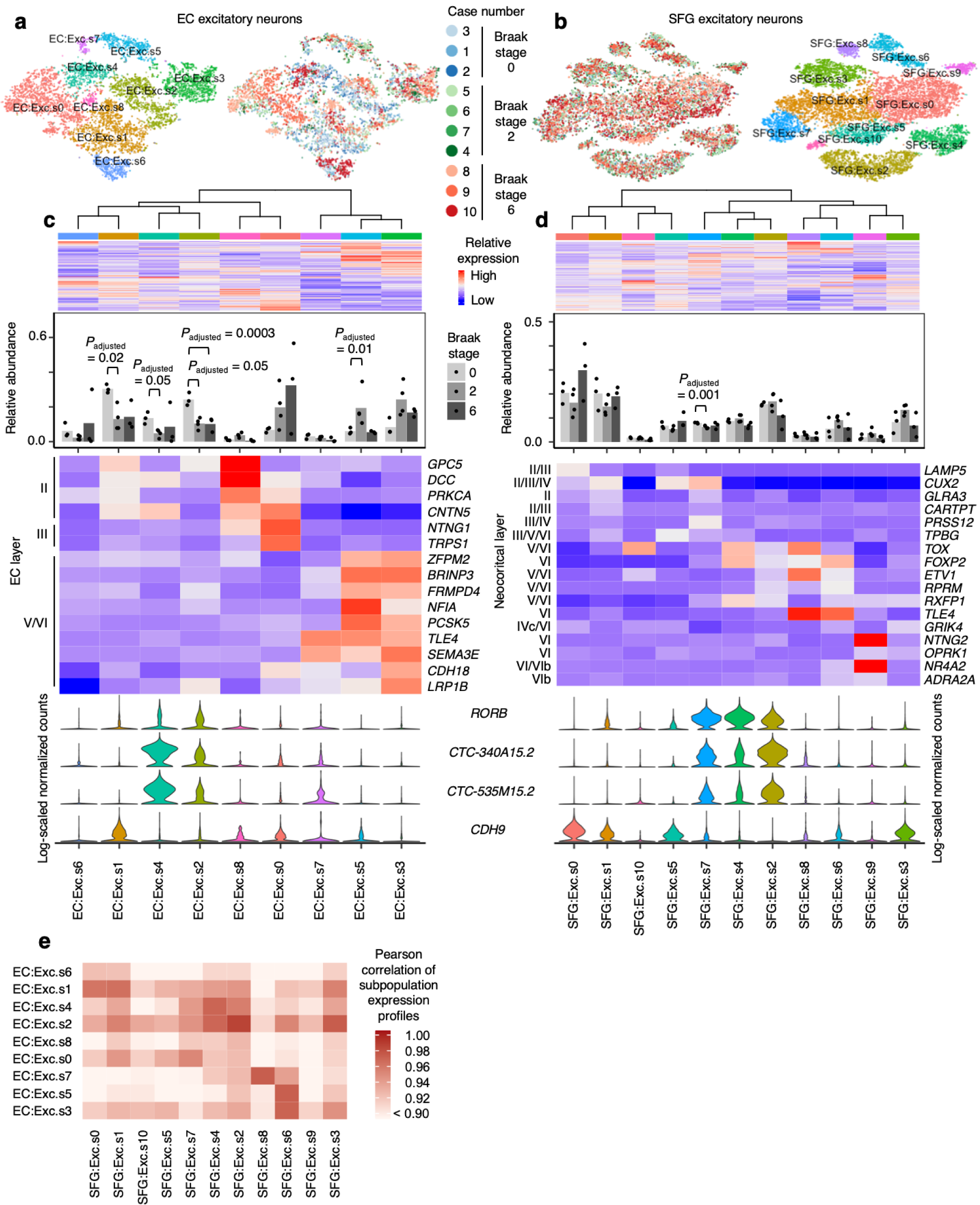
RORB-expressing excitatory neuron subpopulations in the EC are selectively vulnerable. **a-b**, tSNE projection of excitatory neurons from the EC (**a**) and SFG (**b**) in their respective alignment spaces, colored by individual of origin (center) or subpopulation identity (outer). **c-d**, Heatmap and hierarchical clustering of subpopulations and subpopulation marker expression (top subpanel); “High” and “Low” relative expression reflect above- and below-average expression, respectively (see Methods). Relative abundance of subpopulations across Braak stages (second subpanel). Expression heatmap of EC layer-specific genes identified from Ramsden *et al*.^34^ (**c**, third subpanel). Expression heatmap of neocortical layer-specific genes from Lake *et al*.^19^ (**d**, third subpanel). Expression of selectively vulnerable subpopulation markers identified in the EC (bottom subpanel). **e**, Heatmap of Pearson correlation between the gene expression profiles of EC and SFG subpopulations.

To identify molecular markers of selectively vulnerable excitatory neuron subpopulations in the EC (EC:Exc.s2, EC:Exc.s4, EC:Exc.s1), we inspected transcript levels of genes differentially expressed between pairs of subpopulations and curated a set of genes which were specifically expressed by no more than four subpopulations (Extended Data Fig. 2a), which we decided was a reasonable threshold for a positive marker to be useful. We found that EC:Exc.s2 and EC:Exc.s4 specifically expressed *RORB, CTC-340A15*.*2* and *CTC-535M15*.*2* (Fig. 2c). *RORB* (RAR-related Orphan Receptor B) encodes a transcription factor known as a marker and developmental driver of layer IV neurons in the neocortex^37-39^, but is also expressed by neurons in other layers^20^. Little is known about the non-coding transcripts *CTC-340A15*.*2* and *CTC-535M15*.*2* in the context of neuronal identity and function. We also found that EC:Exc.s1 was marked by high expression of *CDH9* (Fig. 2c), a cadherin with neuron-specific expression. However, *CDH9* was also expressed by other excitatory neuron subpopulations in the EC, and we could not find markers that were specifically expressed only in EC:Exc.s1. Therefore, we chose to focus our analysis on EC:Exc.s2 and EC:Exc.s4.

In addition to identifying molecular markers of the selectively vulnerable EC:Exc.s2 and EC:Exc.s4 neurons, we also enumerated genes that were differentially expressed in EC:Exc.s2 and EC:Exc.s4 compared to all other excitatory neurons in the EC, controlling for differences across individuals (see Methods). We found that genes with higher expression in EC:Exc.s2 and EC:Exc.s4 were enriched for axon-localized proteins and voltage-gated potassium channels, whereas genes with lower expression in EC:Exc.s2 and EC:Exc.s4 were enriched for synapse-and dendrite-localized proteins and pathways involving G-protein mediated signaling, ion transport, and neurotransmitter receptor signaling (Extended Data Fig. 2b-e, Supplementary Table 1).

We also performed differential gene expression analysis across Braak stages for EC excitatory neuron subpopulations (see Methods), choosing to focus on comparing Braak stage 6 vs. 0, which yielded the largest number of differentially expressed genes. We found a broad decrease in expression of genes encoding pre- and post-synaptic proteins in Braak stage 6 vs. 0 for many EC excitatory neuron subpopulations (Extended Data Fig. 3b,d,f). Furthermore, we observed that EC:Exc.s2, which demonstrated a statistically significant decline in relative abundance in Braak stage 6 vs. 0, also had the largest number of downwardly differentially expressed genes and the strongest enrichments for pre- and post-synaptic proteins in these genes (Extended Data Fig. 3b,d). Overall, the downregulation of synapse-related genes we have observed mirrors the findings from a recent preprint by Marinaro *et al*.^*40*^, which examined the frontal cortex in familial monogenic AD using snRNA-seq, and is consistent with a previous study of gene expression changes in AD in the entorhinal cortex and other brain regions employing laser capture microdissection of neurons followed by DNA microarray analysis^17^.

Having identified and characterized selectively vulnerable excitatory neuron subpopulations in the EC, we next examined excitatory neuron subpopulations in the SFG. Similar to previous studies^19, 20^, we found that excitatory neuron subpopulations in the SFG (11 in total) expressed distinct sets of neocortical layer-specific genes (Fig. 2b,d), recapitulating the laminar organization of the neocortex. Interestingly, SFG:Exc.s4 and SFG:Exc.s2, which were marked by *RORB, CTC-340A15*.*2* and *CTC-535M15*.*2*, trended towards decreased relative abundance only in Braak stage 6 (Fig. 2d; SFG:Exc.s4 *P*_unadjusted_ = 0.06, SFG:Exc.s2 *P*_unadjusted_ = 0.36), consistent with the late appearance of neurofibrillary inclusions in the SFG starting at Braak stage 5. On the other hand, SFG:Exc.s7, which was also marked by *RORB, CTC-340A15*.*2* and *CTC-535M15*.*2*, exhibited a statistically significant but small decrease in relative abundance in Braak stage 2; however, we did not interpret this as a sign of tau pathology-associated selective vulnerability given that neurofibrillary inclusions should not be present in the SFG in Braak stage 2.

Given that SFG:Exc.s4 and SFG:Exc.s2 expressed similar markers as EC:Exc.s4 and EC:Exc.s2, we wondered if SFG:Exc.s4 and SFG:Exc.s2 may resemble EC:Exc.s4 and EC:Exc.s2 more broadly at the transcriptome level. To test this, we calculated the Pearson correlation coefficient between the expression profiles of SFG and EC subpopulations and found that SFG:Exc.s4 and SFG:Exc.s2 were indeed most similar to EC:Exc.s4 and EC:Exc.s2 (Fig. 2e). This finding is consistent with the reported similarity between deep layer neocortical excitatory neurons and EC excitatory neurons in general^41^. Furthermore, this correspondence was preserved when we mapped subpopulations in the EC to those in the SFG by performing cross-sample alignment for both brain regions jointly (Extended Data Fig. 4). The similarity in transcriptomes of vulnerable excitatory neurons in different brain regions is intriguing and suggests similar mechanisms of selective vulnerability in different brain regions.

Although the decrease in the relative abundance of SFG:Exc.s2 and SFG:Exc.s4 in Braak stage 6 was not statistically significant after correction for multiple testing, we asked if we could detect signs of selective vulnerability in neocortical *RORB*-expressing excitatory neurons in an independent dataset with a larger sample size. To this end, we reanalyzed data from Mathys *et al*.^21^, which profiled the prefrontal cortex from 24 AD cases and 24 healthy controls, with our cross-sample alignment pipeline and performed subclustering of excitatory neurons. In the Mathys *et al*. dataset^21^, we discerned 10 excitatory neuron subpopulations, each of which expressed distinct sets of neocortical layer-specific genes (Extended Data Fig. 5a,b) similar to Lake *et al*.^19^ and our dataset. Of these 10 subpopulations, Mathys:Exc.s4, Mathys:Exc.s5, and Mathys:Exc.s1 expressed *RORB* at high levels (*CTC-340A15*.*2* and *CTC-535M15*.*2* were not available in the pre-processed Mathys *et al*.^21^ data). Importantly, we observed a statistically significant decrease in the relative abundance of Mathys:Exc.s4 in male AD cases vs. controls (Extended Data Fig. 5b), recapitulating the selective vulnerability observed in our dataset, which consists only of male individuals. Furthermore, gene expression correlation analysis showed that Mathys:Exc.s4 was the most similar to EC:Exc.s2 and EC:Exc.s4 (Extended Data Fig. 5c), again demonstrating similarity between selectively vulnerable excitatory neurons in the neocortex and those in the EC.

Although we did not detect any statistically significant changes in the relative abundance of *RORB*-expressing subpopulations in female individuals in Mathys *et al*.^21^, Mathys.Exc.s1 trended towards decreased relative abundance in female AD cases (*P*_unadjusted_ = 0.17) and mapped to EC:Exc.s2 by gene expression correlation (Extended Data Fig. 5b,c). Furthermore, Marinaro *et al*.^40^ included both male and female cases of monogenic AD and also reported the selective vulnerability of two out of four *RORB*-expressing excitatory neuron subpopulations in the prefrontal cortex (ExcB1 and ExcB4)^40^, providing further evidence that subsets of *RORB*-expressing excitatory neurons in the neocortex are selectively vulnerable.

Considering the Mathys *et al*.^21^ and the Marinaro *et al*.^40^ datasets together with our dataset, it appears that while not all *RORB*-expressing excitatory neuron subpopulations in the neocortex showed signs of selective vulnerability, those that did were the most similar to *RORB*-expressing excitatory neurons in the EC, all of which showed signs of selective vulnerability.

### Validation of the selective vulnerability of *RORB*-expressing excitatory neurons

To validate our finding from snRNA-seq that *RORB*-expressing excitatory neurons in the EC were highly vulnerable in AD, we performed multiplex immunofluorescence on post-mortem samples from a larger cohort of individuals (Table 1). Specifically, we quantified the proportion of excitatory neurons and RORB-positive excitatory neurons in the EC superficial layers (i.e. above layer IV, which we also refer to as dissecans-1^42^ in Fig. 3b) in postmortem tissue from 26 individuals spanning Braak stage 0 to 6, who were devoid of non-AD neuropathological changes (Table 1). Given the heterogeneity of the EC, the areas selected for analysis in the caudal EC were delimited using rigorous cytoarchitectonic parameters to minimize the odds of artifactual results (Fig. 3a-c, Extended Data Fig. 6, see Methods). We used multiplex immunofluorescence^43^ to label cells (DAPI), excitatory neurons (TBR1), RORB+ neurons, and phospho-tau neuronal inclusions (CP-13, Ser 202). We failed to find statistically significant changes in the proportion of excitatory neurons overall (TBR1+ cells among all cells) across disease progression (Fig. 3d). However, we observed a substantial reduction in the proportion of RORB+ neurons among excitatory neurons in Braak stages 2-4 and 5-6 compared to Braak stages 0-1 (Fig. 3e). Furthermore, by analyzing a subset of cases, we detected phospho-tau (CP-13) preferentially in RORB+ compared to RORB-excitatory neurons (Fig. 3f-g). Thus, the above results substantiate that RORB-expressing excitatory neurons are highly vulnerable in AD and that their depletion parallels the accumulation of tau neurofibrillary inclusions.

**Fig. 3.**
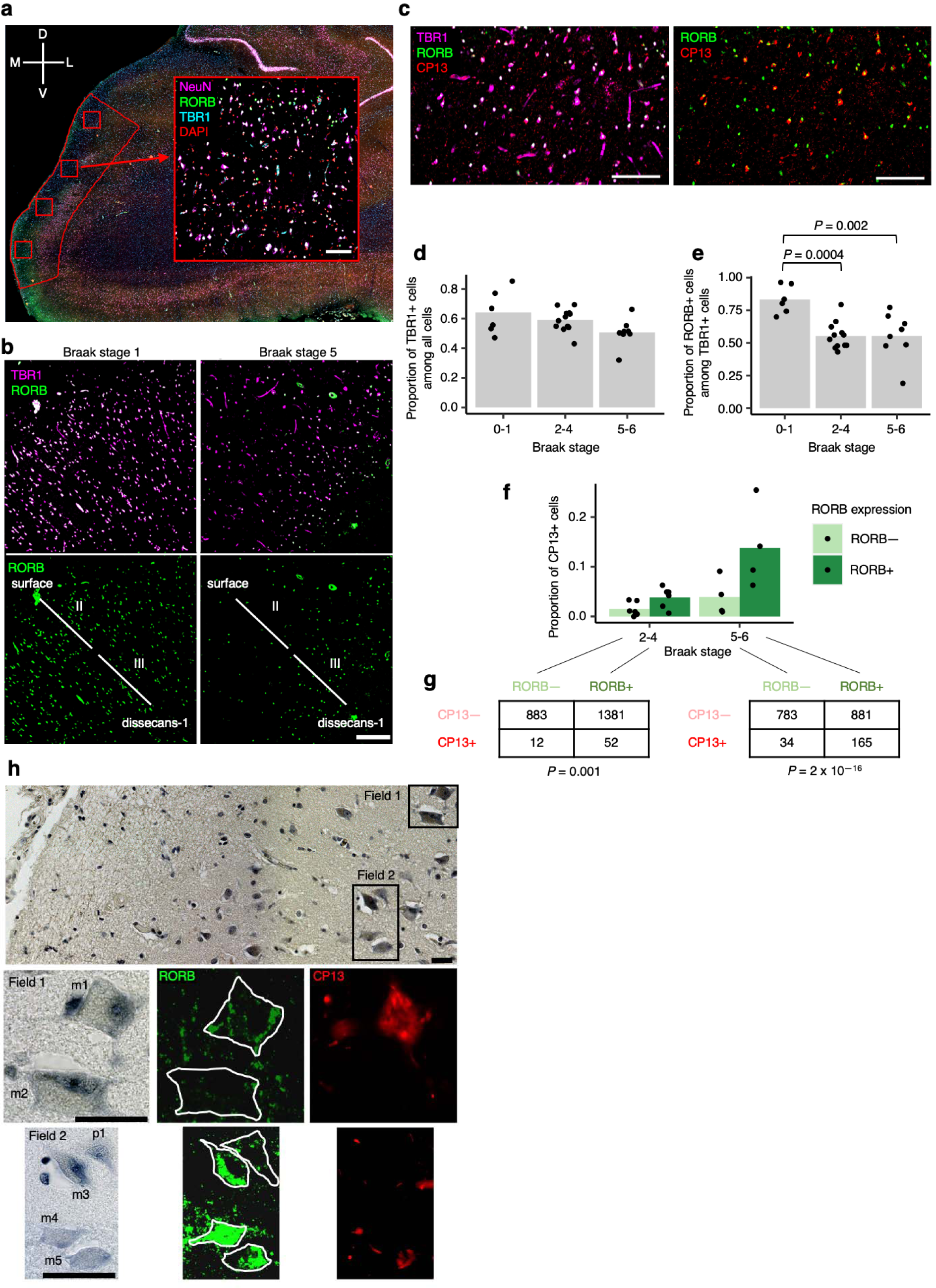
Immunofluorescence of the EC validates selective vulnerability of RORB-expressing excitatory neurons. **a**, The method for extracting regions of interest (ROI) is illustrated using a representative brain slice used for immunofluorescence with the EC delineated in red. Four ROIs (drawn in red squares) were randomly distributed along the superficial layers of the EC and extracted for quantification after masking neurons (see Methods). A representative ROI image with DAPI, NeuN, TBR1, and RORB staining is shown. The anatomical orientation of the slice is provided in the top left corner (D – dorsal, V – ventral, M – medial, L – lateral). **b**, Representative RORB staining in a Braak stage 1 sample (left) vs. a Braak stage 5 sample (right), shown with (top) and without (bottom) excitatory neurons marked by TBR1 staining. The EC layers captured in the image are demarcated in the bottom subpanels (see Methods and Extended Data Fig. 6). **c**, Representative CP13 staining in a Braak stage 6 sample, shown together with TBR1 and RORB staining (left) or only with RORB staining (right). **d-e**, Proportion of TBR1+ cells among all cells (**d**) or proportion of RORB+ cells among TBR1+ cells (**e**) averaged across ROIs for each individual across groups of Braak stages. **f**, Proportion of CP13+ cells in RORB- or RORB+ excitatory neurons (i.e. TBR1+ cells) averaged across ROIs for each individual across groups of Braak stages. **g**, Contingency tables of raw counts of TBR1+ cells based on their RORB or CP13 staining status summed across ROIs and individuals for each group of Braak stages; the Fisher’s Exact Test p-value is shown below each table. **h**, Representative image of EC layer II neurons stained with gallocyanin (top subpanel) with the corresponding RORB and CP13 immunofluorescence signal shown in selected fields (Field 1 – middle subpanels, Field 2 – bottom subpanels). RORB+ neurons include both large multipolar neurons (m1, m3, m4, m5) and pyramidal neurons (p1). One large multipolar neuron (m2) is RORB-. The neuronal somas are outlined manually in white in the RORB immunofluorescence images to aid interpretation. Scale bars shown in **a-c** correspond to 100 microns; scale bars shown in **h** correspond to 15 microns.

Given that large multipolar neurons of “stellate” morphology in EC layer II have been known to be particularly vulnerable in AD^7-10^, we next examined RORB+ excitatory neurons more closely in terms of morphology by overlaying immunofluorescence with Nissl staining. We found that RORB+ excitatory neurons adopted various shapes, including both pyramidal and multipolar morphologies (Fig. 3h). Conversely, some large multipolar neurons are RORB-negative (Fig. 3h). Thus, our results are consistent with the known vulnerability of large multipolar EC layer II neurons, but also demonstrate that a molecular characterization of vulnerable neurons refines the results of morphological studies.

### Lack of differences in vulnerability of inhibitory neuron subpopulations

Having validated the selective vulnerability of a subpopulation of excitatory neurons, we proceeded to examine inhibitory neurons. It has previously been reported that inhibitory neurons are more resistant to tau pathology compared to excitatory neurons in AD^15, 31^. To investigate whether there are differences among inhibitory neuron subtypes in resilience, we performed subclustering of inhibitory neurons in our dataset, discerning 11 subpopulations in the EC and 10 subpopulations in the SFG (Fig. 4a-d). In both brain regions, inhibitory neuron subpopulations expressed distinct sets of inhibitory neuron subtype markers (Fig. 4a-d), consistent with previous studies^19, 20^. We did not any detect statistically significant changes in the relative abundance of inhibitory neurons subpopulations in the EC or SFG (Fig. 4c-d), or in the prefrontal cortex in Mathys *et al*.^21^ (Extended Data Fig. 7). Although Marinaro *et al*. reported broad depletion of inhibitory neuron subpopulations in familial monogenic AD, there was no strong evidence of *selective* vulnerability in particular inhibitory neuron subpopulations relative to other inhibitory neuron subpopulations in Marinaro *et al*.

**Fig. 4.**
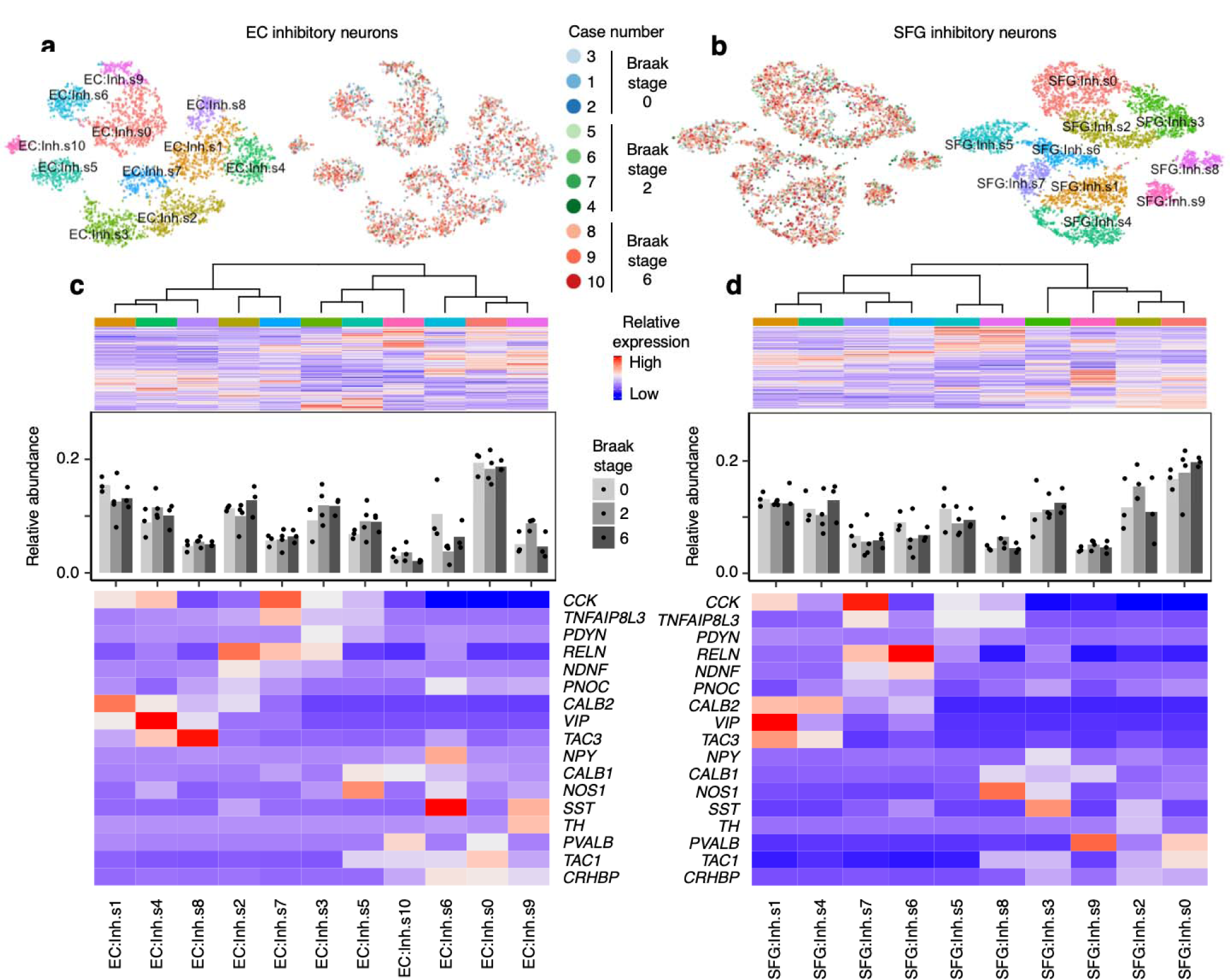
Inhibitory neuron subpopulations do not consistently show differences in resilience or vulnerability to AD progression. **a-b**, tSNE projection of inhibitory neurons from the EC (**a**) and SFG (**b**) in their respective alignment spaces, colored by individual of origin (center) or subpopulation identity (outer). **c-d**, Heatmap and hierarchical clustering of subpopulations and subpopulation marker expression (top subpanel); “High” and “Low” relative expression reflect above- and below-average expression, respectively (see Methods). Relative abundance of subpopulations across Braak stages (middle subpanel). Expression heatmap of inhibitory neuron molecular subtype markers from Lake *et al*.^19^ (bottom subpanel).

### Analysis of glial subpopulations

Glial cells have emerged as important players in AD. We found a trend towards increased relative abundance of microglia in the EC in with AD progression (Fig. 1f), consistent with microgliosis. Next, we asked whether a specific transcriptional state of microglia is associated with AD in our dataset. Recent single-cell profiling of microglia from mouse models of AD identified disease-associated microglia^44^ (DAM), the transcriptional signature of which overlap only partially with that of human microglia found in AD^45^. Considering the possibility that DAMs may cluster separately from homeostatic microglia after cross-sample alignment, we performed subclustering of microglia in our dataset, discerning 4 subpopulations in the EC and 5 subpopulations in the SFG (Extended Data Fig. 8a-b). However, similar to Thrupp *et al*.^46^, we were unable to detect the expression of the majority of homeostatic microglia markers and DAM markers in our dataset or in Mathys *et al*.^21^ (Extended Data Fig. 8d-f), which may be due to the relatively low number of genes captured in microglia compared to other cell types (Fig. 1h-i) and the depletion of many DAM markers in nuclei compared to whole cells^46^.

We next examined oligodendrocytes, which have been shown by Mathys *et al*.^21^ to exhibit a strong transcriptional response in AD. Subclustering of oligodendrocytes in the EC and SFG revealed subpopulations (EC:Oligo.s0 and EC:Oligo.s4, SFG:Oligo.s1 and SFG:Oligo.s2) which exhibited higher expression of AD-associated oligodendrocyte genes from Mathys *et al*.^21^, i.e. genes with higher expression in the AD-associated subpopulation Oli0 in Mathys *et al*.^21^ (Extended Data Fig. 9d-e). Although the function of these genes in the context of AD is largely unknown, a spatial transcriptomics study of AD^47^ has recently implicated a subset of these genes in the response of oligodendrocytes to amyloid plaques (e.g. *CRYAB, QDPR*).

Lastly we turned our attention to astrocytes. While reactive astrocytes are ubiquitously associated with AD pathology^48, 49^, only few studies to date have directly profiled reactive astrocytes due to the difficulty of specifically isolating reactive astrocytes^50, 51^. Similarly to our interrogation of microglia, we asked if reactive astrocytes would cluster separately from non-reactive astrocytes after cross-sample alignment. After subclustering of astrocytes in our dataset, we discerned 4 subpopulations in the EC and 6 subpopulations in the SFG (Fig. 5a-d). In each brain region, there was at least one subpopulation (EC:Astro.3, SFG:Astro.s4 and SFG:Astro.s5) that expressed dramatically higher levels of *GFAP*, which we will refer to as *GFAP*_high_ astrocytes (Fig. 5c,d). In the EC, *GFAP*_high_ astrocytes also expressed *CD44* and *HSPB1*, markers of pan-reactive astrocytes^52^; *TNC*, which is upregulated in stab-wound reactive astrocytes^53, 54^; and *HSP90AA1*, which is upregulated in reactive astrocytes associated with middle cerebral artery occlusion^55^ (Fig. 5c,d). Interestingly, in the SFG, *GFAP*_high_ astrocytes consisted of two subpopulations, one marked by higher expression of *CD44* and *TNC*, both of which are involved in interactions with the extracellular matrix, and the other marked by higher expression of *HSPB1* and *HSP90AA1*, both of which are chaperones involved in proteostasis. In terms of downregulated genes, *GFAP*_high_ astrocytes consistently expressed lower levels of genes associated with glutamate/GABA homeostasis (*SLC1A2, SLC1A3, GLUL, SLC6A11*; see Methods for references) and synaptic adhesion/maintenance (*NRXN1, CADM2, PTN, GPC5*; see Methods for references), suggesting a loss of homeostatic function.

**Fig. 5.**
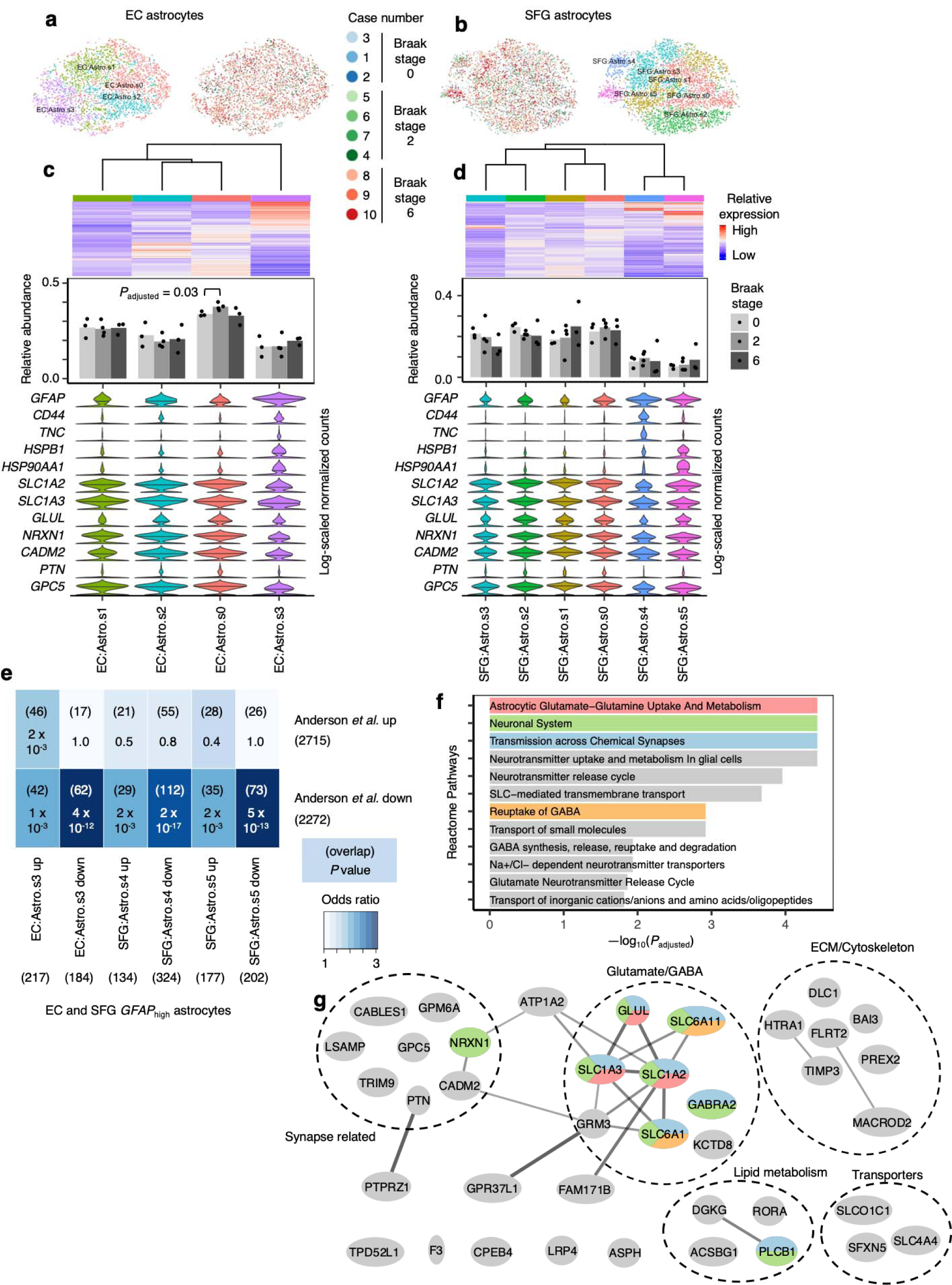
*GFAP*_high_ astrocytes show signs of dysfunction in glutamate homeostasis and synaptic support. **a-b**, tSNE projection of astrocytes from the EC (**a**) and SFG (**b**) in their respective alignment spaces, colored by individual of origin (center) or subpopulation identity (outer). **c-d**, Heatmap and hierarchical clustering of subpopulations and subpopulation marker expression (top subpanel); “High” and “Low” relative expression reflect above- and below-average expression, respectively (see Methods). Relative abundance of subpopulations across Braak stages (middle subpanel). Expression of genes associated with reactive astrocytes, with median expression level marked by line (bottom subpanel). **e**, Enrichment analysis of overlap between differentially expressed genes in *GFAP*_high_ astrocytes vs. differentially expressed genes in reactive astrocytes from Anderson *et al*.^56^ The number of genes in each gene set and the number of overlapping genes are shown in parentheses, and the hypergeometric test p-values (corrected for multiple testing using the Benjamini-Hochberg procedure) are shown without parentheses. **f**, Enrichment of Reactome pathways in downregulated genes in *GFAP*_high_ astrocytes, with selected terms highlighted in color. **g**, Functional association network (see Methods) of downregulated genes shared between EC and SFG *GFAP*_high_ astrocytes that overlap with those in Anderson *et al*.^56^ Genes with stronger associations are connected by thicker lines. Genes that belong to selected gene sets in **k** are highlighted in color.

Examination of all differentially expressed genes in *GFAP*_high_ astrocytes compared to other astrocyte subpopulations showed significant overlap with differentially expressed genes from reactive astrocytes in a mouse model of spinal cord injury^56^ (Fig. 5e). Overlapping downregulated genes included the previously noted genes associated with glutamate homeostasis and synaptic adhesion/maintenance and also genes related to lipid metabolism, cytoskeleton and extracellular matrix, and transporters (Fig. 5f-g).

Finally, to confirm the presence of *GFAP*_high_ astrocytes in an independent dataset, we performed subclustering of astrocytes from Mathys et *al*.^21^ after cross-sample alignment, which yielded 3 subpopulations (Extended Data Fig. 10a,b). Indeed, we found that Mathys:Astro.s2 behaved identically compared to *GFAP*_high_ astrocytes from the EC and SFG in terms of upregulating reactive astrocyte markers and downregulating genes associated with glutamate/GABA homeostasis and synaptic adhesion (Extended Data Fig. 10b). Furthermore, the differentially expressed genes in Mathys:Astro.s3 overlapped highly with those in *GFAP*_high_ astrocytes from the EC and SFG (Extended Data Fig. 10c).

## DISCUSSION

Selective vulnerability is recognized as a fundamental feature of neurodegenerative diseases, including AD. Past studies have characterized the most vulnerable neurons in AD based on topography and morphology. For instance, EC layer II neurons have been found to be more vulnerable than EC layer III pyramidal neurons^10-12^. However, the molecular signature of selectively vulnerable neurons in AD is largely unknown. In this study, we performed snRNA-seq of well-characterized postmortem brain tissue from individuals spanning the progression of AD-type tau neurofibrillary pathology, followed by cross-sample data alignment to identify and characterize selectively vulnerable neuronal populations in the caudal EC and the SFG (Brodmann area 8), representing areas that develop tau neurofibrillary inclusions early and late in the course of AD, respectively. We then validated the snRNA-seq results using quantitative neuropathological methods in a larger cohort spanning all Braak stages of neurofibrillary pathology.

The transentorhinal region and the EC, hubs for integrating information from hippocampal, cortical and subcortical regions^35^, are the first cortical fields to accumulate tau-positive neurofibrillary inclusions followed by neuronal loss in AD^3^. The transentorinal region is first affected in Braak stage 1, followed by the EC in Braak stage 2. The EC is a relatively phylogenetically conserved brain structure in mammals^35, 57^. The rodent EC can be subdivided into medial and lateral portions based on cytoarchitectonics and projections. In primates, the EC has been subdivided into up to 16 regions that show differential abundances of several neuronal markers, distinct projections, and variation of laminar features^11, 42, 57^. During evolution, the position of the EC changed, and the mouse medial EC (the source of our layer-specific marker genes) is generally regarded as the equivalent of the caudal EC in humans (our sampling location)^36^. Irrespective of the parcellation scheme adopted, the EC is a heterogeneous structure and cytoarchitectonic considerations matter when analyzing and sampling this region to avoid biased observations^42^.

Neurons in EC layer II are particularly vulnerable in AD^7, 9, 10, 58^. EC layer II features a mixture of neuronal subpopulations defined by morphology. Large multipolar neurons (“stellate cells”), which are deemed to be excitatory, are abundant^59^. Other neurons assume smaller multipolar morphology and variably sized pyramidal, bipolar, spindle-shaped, and triangular morphologies. Large multipolar neurons are prone to develop AD-tau inclusions and degenerate in very early AD stages^6, 60^. However, variation in the size, shape, and density of these large multipolar neurons along mediolateral and rostrocaudal gradients^42, 60^ has hampered rigorous quantitative characterization of their depletion in AD.

Here, we discovered that in the caudal EC, specific excitatory neuron subpopulations defined by snRNA-seq were selectively vulnerable in AD, exhibiting a ∼50% decline in their relative abundance already in early AD stages. These neurons expressed genes associated with layer II of the mouse medial EC, consistent with the known vulnerability of neurons in the superficial layers of the human EC in AD^7-10^.

Importantly, we identified *RORB* as a marker of these selectively vulnerable excitatory neuron subpopulations, and subsequently validated the selective depletion of RORB+ excitatory neurons in the EC along AD progression by counting these neurons in a larger cohort of individuals using multiplex immunofluorescence. We found that the selectively vulnerable RORB+ excitatory neurons included both large multipolar neurons and pyramidal neurons. Although this finding is consistent with the known vulnerability of large multipolar EC layer II neurons, it also demonstrates that morphology alone is insufficient to determine gradients of selective vulnerability.

We then showed that tau neuronal inclusions, a chief AD neuropathological hallmark, preferentially accumulated in RORB+ excitatory neurons in the EC. To uncover potential cell biological mechanisms underlying the vulnerability of EC RORB+ excitatory neurons, we compared the gene expression profiles of EC *RORB*-expressing excitatory neurons against all other EC excitatory neurons, which revealed differences in the expression of genes encoding synapse-vs. axon-localized proteins, potassium channel subunits, G-protein signaling molecules, and neurotransmitter receptor signaling molecules. Future studies utilizing *in vitro* and animal models of AD together with techniques for manipulating gene expression such as CRISPR inhibition and activation^61-63^ will make it possible to address these potential mechanistic connections among *RORB*-expression, phospho-tau accumulation, and vulnerability.

In neocortical areas, layers III and V are the first to accumulate tau neurofibrillary inclusions in AD^3, 32, 33^. We found that in the SFG, a subset of *RORB*-expressing excitatory neuron subpopulations showed signs of selective vulnerability only late in AD, in line with the late appearance of neurofibrillary inclusions in the SFG, although the decrease in their relative abundance did not pass our threshold for statistical significance after correction for multiple testing. Interestingly, we found through correlation analysis of gene expression and also EC-SFG cross-sample alignment that *RORB*-expressing excitatory neuron subpopulations in the SFG showing signs of selective vulnerability were similar to those in the EC in terms of their transcriptomic profile. To verify the reproducibility of our findings, we re-analyzed the data from Mathys *et al*.^21^ using our cross-sample alignment approach. Although Mathys *et al*.^21^ probed a different neocortical region (the prefrontal cortex), we found that one of their *RORB*-expressing excitatory neuron subpopulations also exhibited selective vulnerability and mapped to our *RORB*-expressing excitatory neuron subpopulations in the EC. In addition, Marinaro *et al*.^*40*^, which examined familial monogenic AD, also reported the selective depletion of two out four *RORB-*expressing excitatory neuron subpopulations in their dataset. Considering our dataset jointly with the Mathys *et al*.^21^ and Marinaro *et al*.^*40*^ datasets, it appears that in the neocortex, while not all *RORB*-expressing excitatory neuron subpopulations are selectively vulnerable, those that are vulnerable have a similar transcriptional profile as selectively vulnerable neurons in the EC. Given that RORB is known to function as a developmental driver of neuronal subtype identity in the neocortex^37-39^, we hypothesize that the vulnerability of *RORB*-expressing excitatory neuron subpopulations in different brain regions is caused by the activity of RORB and potentially other subtype-determining transcription factors, which drive gene expression programs that confer an intrinsic vulnerability of these neurons that is realized in the presence of AD-tau pathology. Further mechanistic studies involving the perturbation of *RORB* expression in animal models of AD are necessary to test this hypothesis.

A previous study suggested changes in the number of neurons expressing calbindin and parvalbumin, which tend to mark inhibitory neurons, in EC layer II in AD^58^. Here, we found no evidence of selective vulnerability in inhibitory neurons subpopulations in EC layer II or any other layer. Inhibitory neurons in the EC superficial layers show a gradient of abundance in the various EC regions^35^, which could confound the results. But, given that we used strict cytoarchitectonic criteria to sample the EC, it is unlikely that our results reflect comparisons of different EC areas across the cases. Also, evidence suggest that these inhibitory neurons undergo changes in morphology and function, rather than loss in sporadic AD^58^. Thus, our results do not preclude the possibility that inhibitory neuron subpopulations may be differentially affected by AD progression at the morphological and likely functional level, even if neuronal loss is not apparent.

Until recently, AD research was mostly neuron-centric, but accumulating evidence is highlighting the importance of glial changes in the pathogenesis of AD. Although we could not detect the disease-associated microglia signature^44, 45^ in our study, likely due the low number of transcripts recovered in microglia, we discovered an astrocyte subpopulation expressing high levels of *GFAP*, which we termed *GFAP*_high_ astrocytes, in both the EC and SFG, as well as in the prefrontal cortex from Mathys *et al*.^21^ We found that *GFAP*_high_ astrocytes also expressed higher levels of other genes associated with reactive astrocytes, while expressing lower levels of genes involved in glutamate homeostasis and synaptic adhesion/maintenance, which suggests loss of normal astrocyte homeostatic functions. Furthermore, we found a high degree of overlap between genes differentially expressed in *GFAP*_high_ astrocytes and genes differentially expressed in reactive astrocytes from a mouse model of spinal injury^56^. Thus, we believe that *GFAP*_high_ astrocytes correspond to reactive astrocytes in AD, which may have compromised homeostatic function.

Our study has several methodological strengths. First, the postmortem cohort used for snRNA-seq and histopathological validation consists of well-characterized cases, devoid of non-AD pathology. To minimize confounders in the snRNA-seq results, we selected only male cases with an *APOE* ε3/ε3 genotype. Second, we sequenced a very large number of nuclei from each case (∼10,000 nuclei per case, compared to ∼1,700 nuclei per case in Mathys *et al*.^21^) from two brain regions per individual (∼4,000 nuclei from the EC and ∼6,000 nuclei from the SFG). Third, the human cortex displays a complex parcellation scheme based on cytoarchitectonic characteristics that reflect differences in the abundance of various cell subpopulations, with implications for function, projections, and differential vulnerability in AD. Many RNA-seq studies of AD used broad descriptions to define the sampled brain areas, making it challenging to understand if they were sampled from the same subfields. We used strict cytoarchitectonic criteria to sample brain regions for snRNA-seq and histopathological validation. Fourth, our focus was on defining cell type subpopulations that showed changes in relative abundance between disease stages, which can reflect important disease processes such as neuronal loss, and to define the genes characteristic of these subpopulations. The way we defined cell type subpopulations independently of disease progression allowed us to compare gene expression between different cell type subpopulations within individuals while controlling for differences among individuals; this is more robust than comparing gene expression in a given subpopulation across groups of individuals, which can be influenced by differences in confounding factors between the groups. Lastly, by validating our findings using a novel multiplex immunofluorescence approach that enables probing a higher number of antibodies simultaneously^43^, we could quantify the relative abundance of excitatory neurons and RORB+ neurons and also demonstrate that RORB+ excitatory neurons were preferentially affected by neurofibrillary inclusions.

A limitation of our study is that we only included male *APOE* ε3/ε3 individuals in the snRNA-seq analysis. We included females and individuals carrying the APOE ε4 allele associated with AD risk in our histopathological validation, but caution should be taken before generalizing our results to these groups. Future studies will provide a systematic analysis of the impact of sex and *APOE* status on selective vulnerability in AD.

In conclusion, our study contributes, to the best of our knowledge, a pioneering characterization of selectively vulnerable neuronal populations in AD using snRNA-seq profiling of paired brain regions from the same individuals, which were all carefully curated AD cases and controls. These results will inform future studies of the mechanistic basis of selective vulnerability in both animal and *in vitro* models, such as human iPSC-derived neurons, in which the deployment of CRISPR inhibition and activation technology enables elucidation of the functional consequences of transcriptomic changes^61, 64^.

## ONLINE METHODS

### Post-mortem cohort

This study was approved by and University of Sao Paulo institutional review board and deemed non-human subject research by the University of California, San Francisco (UCSF). De-identified human postmortem brain tissue was supplied by the Neurodegenerative Disease Brain Bank (NDBB) at UCSF, and the Brazilian BioBank for Aging Studies (BBAS) from the University of Sao Paulo^65^. The NDBB receives brain donations from patients enrolled in the UCSF Memory and Aging Center research programs. The BBAS is population□based and houses a high percentage of pathologically and clinically normal control subjects who are not available in the NDBB. Neuropathological assessments were performed using standardized protocols and followed internationally accepted criteria for neurodegenerative diseases^66-68^. The brain samples used in this study contained a broad burden of AD-type pathology and were selected to be free from non-AD pathology including Lewy body disease, TDP-43 proteinopathies, primary tauopathies, and cerebrovascular changes. Argyrophilic grain disease (AGD) was not an exclusion criterion based on its high prevalence and lack of correlation with significant clinical symptoms^69-71^. In total, the cohort included 10 cases who underwent snRNA-seq, representing Braak stages 0, 2 and 6, all ApoE 3/3, and 26 cases who underwent neuroanatomical analysis, representing Braak stages 0-6^3, 25^, ranging from 2-5 individuals per Braak stage. Table 1 depicts the characteristics of the 31 cases.

### Isolation of nuclei from frozen post-mortem human brain tissue

Isolation of nuclei was performed similarly as previously described^72^. Briefly, frozen brain tissue was dounce homogenized in 5 ml of lysis buffer (0.25 M sucrose, 25 mM KCl, 5 mM MgCl_2_, 20 mM Tricine-KOH, pH 7.8, 1 mM DTT, 0.15mM spermine, 0.5 mM spermidine, 1X protease inhibitor (Sigma, 4693159001), and RNAse Inhibitor (Promega, N2615)). Following initial dounce homogenization, IGEPAL-630 was added to a final concentration of 0.3% and the sample was homogenized with 5 more strokes. The solution was then filtered through a 40 um cell filter and mixed with Optiprep (Sigma, D1556-250ML) to create a 25% Optiprep solution. This solution was then layered onto a 30%/40% Optiprep gradient and centrifuged at 10,000g for 18 minutes using the SW41-Ti rotor. The nuclei were collected at the 30%/40% Optiprep interface.

### Droplet-based single-nucleus RNA-sequencing

Droplet-based single-nucleus RNA-sequencing (snRNA-seq) was performed using the Chromium Single Cell 3′ Reagent Kits v2 from 10X Genomics. Nuclei were resuspended to a concentration of 1000 nuclei/uL in 30% Optiprep solution before loading according to manufacturer’s protocol, with 10,000 nuclei recovered per sample as the target. cDNA fragment analysis was performed using the Agilent 4200 TapeStation System. Sequencing parameters and quality control were performed as described by The Tabula Muris Consortium^73^.

### Pre-processing of snRNA-seq data

Sequencing data generated from snRNA-seq libraries were demultiplexed using *Cellranger* (version 2.1.0) *cellranger mkfastq*. To align reads, we first generated our own pre-mRNA GRCh38 reference genome using *cellranger mkref* in order to account for introns that may be eliminated using the default GRCh38 reference genome. Alignment and gene expression quantification was then performed using *cellranger count* with default settings.

### Exploratory analysis of EC and SFG data

For each sample, the raw gene-barcode matrix outputted by *Cellranger* (version 2.1.0) was converted into a *SingleCellExperiment* (SCE) object using the *read10xCounts* function from the *DropletUtils* package^74^ (version 1.2.2). Droplets containing nuclei were then distinguished from empty droplets using *DropletUtils::emptyDrops* with the parameter *FDR = 0*.*01*, and then nuclei (hereon also referred to as “cells”) with less than 200 UMIs were discarded. Afterwards, SCE objects corresponding to each sample were merged into a single SCE object for downstream processing and analyses.

For normalization of raw counts, to avoid artifacts caused by data sparsity, the approach of Lun *et al*.^75^ was adopted: For each sample, cells were first clustered using a graph-based method followed by pooling counts across cells in each cluster to obtain pool-based size factors, which were then deconvoluted to yield cell-based size factors. Clustering was performed using the *quickCluster* function from the *scran* package^76^ (version 1.10.2) with the parameters *method = ‘igraph’, min*.*mean = 0*.*1, irlba*.*args = list(maxit = 1000)*, and the *block* parameter set to a character vector containing the sample identity of each cell. Size factors were computed using *scran::computeSumFactors* with the parameter *min*.*mean = 0*.*1* and the *cluster* parameter set to a character vector containing the cluster identity of each cell; cells with negative size factors were removed. Normalization followed by log-transformation was then performed using the *normalize* function from the *scater* package^77^ (version 1.10.1).

Prior to dimensionality reduction, highly variable genes were identified for each sample separately using the approach of Lun *et al*.^76^: Each gene’s variance was decomposed into a technical and biological component. Technical variance was assumed as Poisson and modeled using *scran::makeTechTrend*. The mean-variance trend across genes was fitted using *scran::trendVar* with parameters *use*.*spikes = FALSE* and *loess*.*args = list(span = 0*.*05)*; and the *trend* slot of the resulting fit object was then set to the output of *scran::makeTechTrend*. Biological variance was extracted from the total variance using *scran::decomposeVar* with the above fit object as the input. Finally, highly variable genes that were preserved across samples were identified by combining the variance decompositions with *scran::combineVar*, using Stouffer’s z-score method for meta-analysis (*method = ‘z’*), which assigns more weight to samples with more cells.

For initial data exploration, genes with combined biological variance greater than 0 were used as the feature set for dimensionality reduction by principal component analysis using *scran::parallelPCA*, which uses Horn’s parallel analysis to decide how many principal components to retain, with parameter *approx = TRUE*. Clustering was then performed on the retained principal components using the *FindClusters* function from the *Seurat* package^78^ (version 2.3.4) with parameter *resolution = 0*.*8*, which required conversion of SCE objects to Seurat objects using *Seurat::Convert*. To visualize the clusters, t-stochastic neighborhood embedding (tSNE) was performed on the retained principal components using *scater::runTSNE* with parameters *perplexity = 30* and *rand_seed = 100*.

### Cross-sample alignment of SFG and EC data

Initial data exploration revealed that clustering was driven by individual of origin in addition to cell type identity, which makes it difficult to analyze changes in the relative abundance or gene expression of a given cell type across disease progression or brain regions. To recover clusters defined by mainly by cell type identity, data was aligned across samples from each brain region using with *scAlign*^29^ (version 1.0.0), which leverages a neural network to learn a low-dimensional alignment space in which cells from different datasets group by biological function independent of technical and experimental factors. As noted by Johansen & Quon^29^, *scAlign* converges faster with little loss of performance when the input data is represented by principal components or canonical correlation vectors. Therefore, prior to running *scAlign*, the top 2000 genes with the highest combined biological variance were used as the feature set for canonical correlation analysis (CCA), which was implemented using *Seurat::RunMultiCCA* with parameter *num*.*cc = 15*. The number of canonical coordinates to use for *scAlign* was determined by the elbow method using *Seurat::MetageneBicorPlot. scAlign* was then run on the cell loadings along the top 10 canonical correlation vectors with the parameters *options = scAlignOptions(steps = 10000, log*.*every = 5000, architecture = ‘large’, num*.*dim = 64), encoder*.*data = ‘cca’, supervised = ‘none’, run*.*encoder = TRUE, run*.*decoder = FALSE, log*.*results = TRUE, and device = ‘CPU’*. Clustering was then performed on the full dimensionality of the ouptut from *scAlign* using *Seurat::FindClusters* with parameter *resolution = 0*.*8* for the SFG and *resolution = 0*.*6* for the EC. Clusters were visualized with tSNE using *Seurat::RunTSNE* on the full dimensinality of the output from *scAlign* with parameter *do*.*fast = TRUE*. Alignment using *scAlign* followed by clustering was also performed for all samples from both brain regions jointly.

To assign clusters identified in the aligned subspace generated by *scAlign* to major brain cell types, the following marker genes were used: *SLC17A7* and *CAMK2A* for excitatory neurons, *GAD1* and *GAD2* for inhibitory neurons, *SLC1A2* and *AQP4* for astrocytes, *MBP* and *MOG* for oligodendrocytes, *PDGFRA* and *SOX10* for oligodendrocyte precursor cells (OPCs), *CD74* and *CX3CR1* for microglia/myeloid cells, and *CLDN5* and *FLT1* for endothelial cells. Clusters expressing markers for more than one cell type, most likely reflecting doublets, were removed from downstream analyses.

### Cell type-specific subclustering (subpopulation) analysis

To identify cell type subpopulations, cells from all samples belonging to a given major cell type were extracted for sample-level re-computation of size factors and highly variable genes. CCA was then performed using the top 1000 genes with the highest combined biological variance as the feature set, followed by alignment of the first 10 to 12 canoical coordinates with *scAlign*, with *steps = 2500*. The full dimensionality of the output from *scAlign* was used for subclustering (using *resolution = 0*.*4*) and tSNE. Analyzing cells from each brain region separately, marker genes for subpopulations were identified using *scran::findMarkers* with parameters *direction = ‘up’, pval*.*type = ‘any’, lfc = 0*.*58*, and the *block* parameter set to a character vector corresponding to each cell’s sample identity. Subpopulations that expressed markers for more than one cell type were removed from downstream analyses.

### Identification of differentially expressed genes in cell type subpopulations

To identify genes differentially expressed by a cell type subpopulation compared to all other subpopulations in a way that accounts for true biological replication (i.e. at the level of individuals), UMI counts of cells from the same individual belonging to the subpopulation of interest or all other subpopulations were summed to obtain “pseudo-bulk” samples, which were then analyzed using *edgeR*^79^ (version 3.24.3) following the approach recommended by Amezquita *et al*.^80^ A false-discovery rate cutoff of 0.1 was used.

### Heatmap visualization of relative gene expression across cell types or cell type subpopulations

For heatmaps of relative gene expression across cell types or cell type subpopulations shown in the figures, log-scaled normalized counts of each gene were z-score transformed across all cells and then averaged across cells in each cluster to enhance visualization of differences among clusters. Thus genes with “high” relative expression have above-average expression (positive z-scores) and genes with “low” relative expression have below-average expression (negative z-scores).

### Functional association network analysis and pathway enrichment analysis of differentially expressed genes

Differentially expressed genes were visualized as a functional association network using String-db^81^ (v11), a protein-protein association network based on known physical interactions, functional associations, coexpression, and other metrics, and Cytoscape^82^ (version 3.7.2), a network visualization software. When generating the networks, the String-db association confidence score cutoff set to 0.5, and the network layout was optimized for visualization using the yFiles Organic Layout. For pathway enrichment analysis, enrichments for Gene Ontology terms and Reactome Pathways were also obtained through String-db, using a false-discovery rate cutoff of 0.05.

### Beta regression

For each brain region, to determine the statistical significance of changes in the relative abundance of a given cluster or cell type across disease progression, the relative abundance was computed for each sample and treated as an independent measurement and beta regression^30^ was performed using the *betareg* package^83^ (version 3.1-1), using the formula *relative*.*abundance ∼ braak*.*stage* for both the mean and precision models, and the bias-corrected maximum likelihood estimator (*type = ‘BC’*). The statistical significance of changes in the proportion of TBR1+ cells and RORB+ cells among TBR1+ cells obtained from immunofluorescence validation were assessed similarly as above using beta regression. To correct for multiple hypothesis testing for each family of tests (e.g. testing all cell type subpopulations for a brain region), Holm’s method was used to adjust *P* values obtained from beta regression to control the family-wise type I error rate at 0.05.

### Entorhinal cortex layer-specific genes

Due to the lack of published data on layer-specific genes for the human EC, layer-specific genes in the mouse medial entorhinal cortex (MEC) were obtained from Ramsden *et al*.^34^. (The MEC is the most phylogenetically similar to the human caudal EC^35, 36^ used in this study.) Specifically, genes with expression specific for layer II, III, and V/VI of the mouse MEC according to the S4 Dataset excel spreadsheet in the supplemental information of Ramsden *et al*.^34^ were mapped to human genes, and cross-referenced against genes differentially expressed across EC excitatory neuron subclusters (obtained using *scran::findMarkers* without setting *direction = ‘up’*).

### Re-analysis of the Mathys *et al*. dataset

To re-analyze the data from Mathys *et al*.^21^ using our cross-sample alignment approach, the filtered matrix of UMI counts (“filtered_count_matrix.mtx”) and associated row and column metadata were downloaded from The AMP-AD Knowledge Portal (Synapse ID: syn18485175). Since clinical metadata was not provided in the column metadata, the individual ID (“projid” column in the column metadata) was cross-referenced with the official ROS-MAP clinical metadata (“ROSMAP_Clinical_2019-05_v3.csv”, synapse ID: syn3191087), which was then cross-referenced with the additional metadata provided in Supplementary Table 1 and 3 from Mathys *et al*.^21^ The filtered UMI counts matrix and the associated row and column metadata were then converted to a *SingleCellExperiment* object for analysis. The cell type assignments from Mathys *et al*.^21^ provided in the column metadata were used for subclustering.

### Functional annotation of differentially expressed genes in *GFAP*_high_ astrocytes

We obtained the functional annotation for differentially expressed genes from the GeneCards website^84^ and verified the primary literature references for glutamate/GABA-related genes^85-90^ and synaptic adhesion/maintenance-related genes^91-94^.

### Quantitative histopathological assessment using multiplex immunofluorescence

#### Delineation of the caudal EC

We used archival paraffin blocks from the UCSF/NBDD and BBAS (Table 1). First, we collected blocks sampling the hippocampal formation anterior to the lateral genicular body from the 10 cases used for the snRNAseq and another 30 cases spanning all Braak stages^3^. To determine if the caudal EC region was present, 8µm thick sections of each block underwent hematoxylin and eosin staining (Extended Data Fig. 8A). We took digital images of the stained sections and aligned each one the most approximate section from a large collection of 400 µm thick serial coronal sections of whole-brain hemispheres stained for gallocyanin provided by co-author Heinsen^42, 95^ (Extended Fig Data 8B). We eliminated blocks from five cases used for snRNA-seq and four of the extra cases for lack of caudal EC. Next, again with the aid of the paired gallocyanin sections, we delineated the borders of the caudal EC in each case (Extended Data Fig. 8A).

The EC is considered a peri- or allocortex, depending on the author^11^. EC parcellation and cytoarchitectonic definitions have been a matter of debate, and here, we are adopting the cytoarchitectonic definitions proposed by Heinsen and colleagues^42^, which is based on the examination of thick histological preparations and considered the definitions proposed by Insausti and Amaral (6 layers)^96^ and Braak and Braak (3 layers)^11^. In thick histological sections, the caudal entorhinal region features well-delineated clusters of stellate or principal cells in layer II (pre-alpha clusters) and three lamina dissecans^42^. The external dissecans (dissecans-ext) divides layers II and III is particularly prominent in the caudal EC. Dissecans-1 (diss-1) corresponds to layer IV of Insausti^57^ and the lamina dissecans of Braak and Braak^11^ and Rose^97^. The most internal dissecans (dissecans-2, or diss-2) is hardly appreciated in thin sections but easy to visualize in thick sections. It roughly corresponds to layer Vc of the caudal subregions of Insausti^57^.

#### Multiplex immunofluorescence

Next, for each case, an 8µm thick, formalin-fixed and paraffin-embedded coronal section underwent immunofluorescence against TBR1, RORB and phospho-tau(CP-13) as described below. TBR1, or T-box, brain, 1 is a transcription factor protein that has a role in differentiation of glutamatergic neurons and is a marker for excitatory neurons, including EC excitatory neurons^14, 98^. In summary, sections were deparaffinized and incubated in 3.0% hydrogen peroxide (Fisher, H325-500) in methanol to inactivate endogenous peroxidase. Antigen retrieval was performed in 1X Tris-EDTA HIER solution (TES500) PBS with 0.05% Tween 20 (PBS-T) at pH9 in an autoclave at 121□°C for five□minutes. To reduce nonspecific background staining, sections were blocked with 5% Milk/PBS-T. To avoid cross-reactions between primary antibodies that were raised against the same species, an antibody stripping step using 0.80% β-mercaptoethanol/10% sodium dodecyl sulfate in 12.5% Tris-HCL was performed after the tyramide-signal amplification (TSA) development for RORB.

Sections were first incubated overnight in primary antibody against RORB (1:400, rabbit, HPA008393, Millipore Sigma), which was later developed in goat anti-rabbit HRP (1:400, R-05072-500, Advansta) with Alexa Fluor 488 TSA (1:100, B40953, Thermo Fisher). Next, sections were stripped of RORB primary antibody and then were incubated overnight in a cocktail of primary antibodies against TBR1 (1:100, Rabbit, ab31940, Abcam) and CP13 (1:800, mouse, phospho-tau serine 202, gift of Peter Davies, NY), all of which were later developed with secondary antibodies and fluorophores: for TBR1, Alexa Fluor 546 conjugated anti-rabbit secondary (1:200, A-11010, Thermo Fisher) was used, and for CP13, biotinylated anti-mouse (1:400, BA-2000, Vector Laboratory) with streptavidin Alexa Fluor 790 (1:250, S11378, Thermo Fisher) was used. Sections were then counterstained with DAPI diluted in PBS (1:5000, D1306, Invitrogen). Finally, sections were then incubated in Sudan Black B (199664-25g, Sigma) to reduce autofluorescence and coverslipped using Prolong antifade mounting media (P36980, Invitrogen). A quality control slide was used to verify the efficacy of the antibody stripping process. A detailed description of the method is provided in Ehrenberg et al.^43^ Sections were scanned using a Zeiss AxioScan Slide Scanner.

For generating the images shown in Fig. 3h, a section from case #6 (Braak stage 2, see Table 1) was stained with gallocyanin-chrome alum following standard methods^42^. The section was placed on a cover slip and scanned using a Zeiss AxioScan Slide Scanner. Next, the section was removed from the cover slip and underwent immunofluorescence for RORB and CP13 as described above. Then, the section was placed on a cover slip and scanned once more.

#### Neuronal quantification

The caudal EC delineations carried out in the hematoxylin and eosin-stained slides were then transferred to the immunostained images. Within these borders, we randomly placed four 500×500 µm regions of interest (ROI) overlaying the EC external layers (I to III), which we identified as being external to dissecans-1. We then extracted the ROIs for quantification in ImageJ (Fig. 3). The number of excitatory neurons was quantified by segmenting the TBR1 signal, using a threshold to create a mask and the segmentation editor plugin to manually remove all non-neuronal artifacts and vessels. The number of RORB+ excitatory neurons was then counted using the mask of excitatory (TBR1+) neurons in the segmentation editor and manually removing all neurons not expressing RORB. All segmentations were manually verified for quality control. Quantification was done blinded to the neuropathological diagnosis.We quantified phospho-tau (CP-13) staining in two ROIs in a subset of the cases, using the same FIJI protocol.

## DATA AVAILABILITY

The raw snRNA-seq sequencing data and unfiltered UMI count matrices are available on the Gene Expression Omnibus (GEO) under the accession GSE147528. Single-cell data after quality control is available for download in synapse.org at under the Synapse ID syn21788402. Post quality-control data can also be explored interactively through the CellXGene platform at https://kampmannlab.ucsf.edu/ad-brain.

## CODE AVAILABILITY

We provide the full bioinformatics pipeline for the analysis of snRNA-seq data in this paper at https://kampmannlab.ucsf.edu/ad-brain-analysis.

## ACKNOWLEDGEMENTS

We thank Angela Pisco, Ashley Maynard, Spyros Darmanis and MACA team at the Chan Zuckerberg Biohub for advice on analysis 10X library preparation and reagents. We thank members of the Kampmann lab (Avi Samelson, Xiaoyan Guo, Ruilin Tian, Brendan Rooney) for feedback on the manuscript. This work was supported by NIH awards F30 AG066418 (K.L.), K08 AG052648 (S.S.), R56 AG057528 (M.K., L.T.G.), K24 AG053435 (L.T.G), U54 NS100717 (L.T.G, M.K.), an NDSEG fellowship (E.L.), Alzheimer’s Association fellowship AARF 18-566005 (R.D.R.), FAPESP/CAPES (2016/24326-0) (R.D.R.) and a Chan Zuckerberg Biohub Investigator Award (M.K.). The UCSF Neurodegenerative Disease Brain Bank is supported by NIH grants AG023501 and AG019724, the Tau Consortium, and the Bluefield Project to Cure FTD.

## AUTHOR CONTRIBUTIONS

K.L., E.L., L.T.G. and M.K. conceptualized and led the overall project. K.L., L.T.G. and M.K. wrote the manuscript, with input from all co-authors. K.L. analyzed snRNA-Seq data and visualized results. E.L. generated snRNA-Seq data, with support from R.S., M.T., and N.N. R.D.R., C.K.S., R.E.P.L., C.P. W.W.S., and S.S. contributed to neuropathological data generation and analysis, R.E., A.P. and H.H. contributed to neuropathological data analysis, and S.H.L. contributed to neuropathological method development.

## ETHICS DECLARATIONS

This project was approved the the ethical committee of the University of Sao Paulo (for tissue transfer) and deemed non-human subject research by UCSF.

## EXTENDED DATA

**Extended Data Fig. 1.**
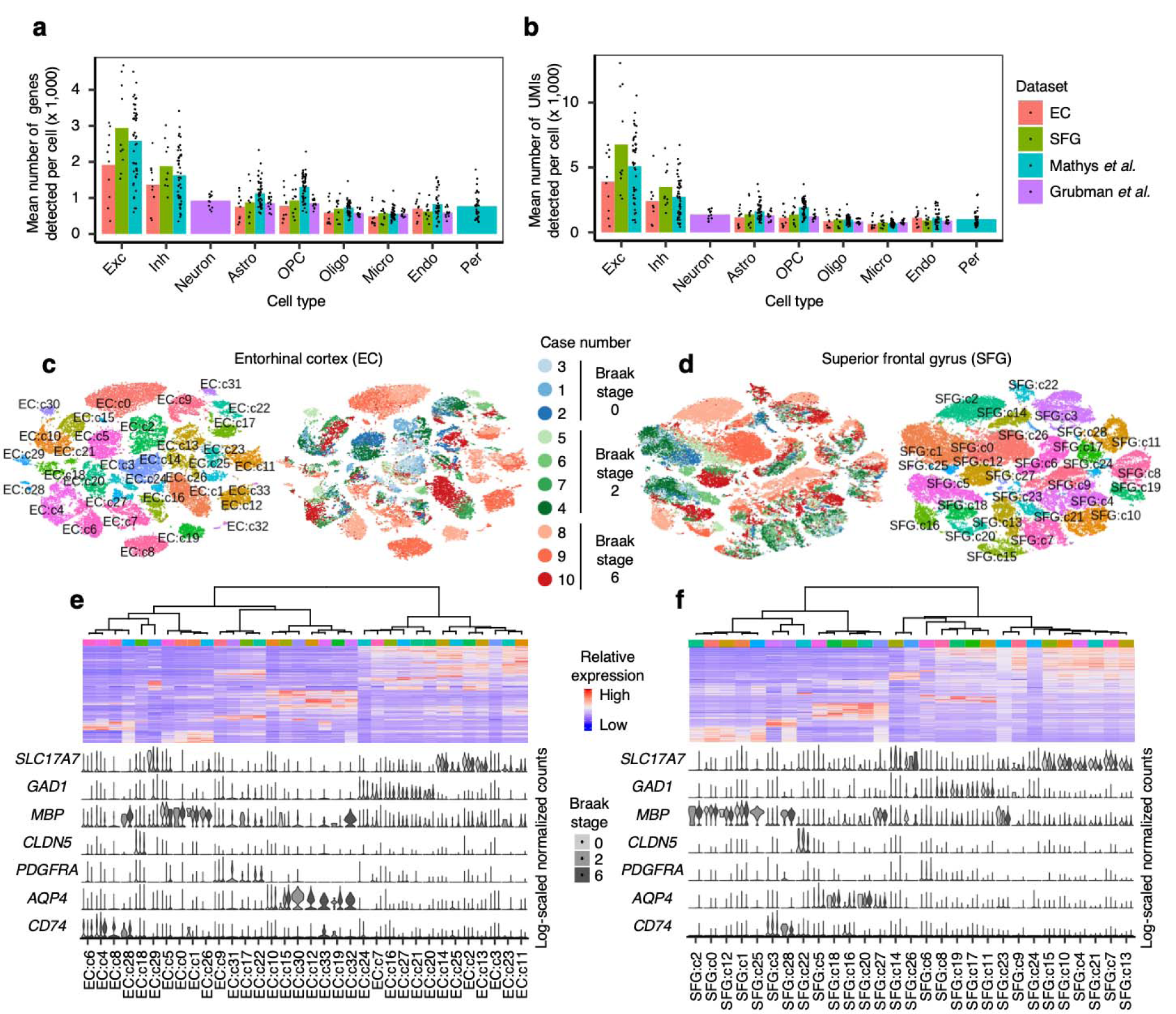
Data quality and initial clustering without cross-sample alignment. **a-b**, Mean number of genes (**a**) or UMIs (**b**) detected per cell across individual samples for major cell types identified in each dataset. Grubman *et al*.^22^ did not resolve excitatory neurons from inhibitory neurons. Pericytes were identified only in Mathys *et al*.^21^ Cell type abbreviations: Exc – excitatory neurons, Oligo – oligodendrocytes, Astro – astrocytes, Inh – inhibitory neurons, OPC – oligodendrocyte precursor cells, Micro – microglia, Endo – endothelial cells, Per – pericytes. **c-d**, tSNE projection of cells from the EC (**c**) and SFG (**d**) clustered without first performing cross-sample alignment, colored by individual of origin (center) or cluster assignment (outer). **e-f**, Heatmap and hierarchical clustering of clusters and cluster marker expression (top subpanels); “High” and “Low” relative expression reflect above- and below-average expression, respectively (see Methods). Expression of cell type markers (bottom subpanels).

**Extended Data Fig. 2.**
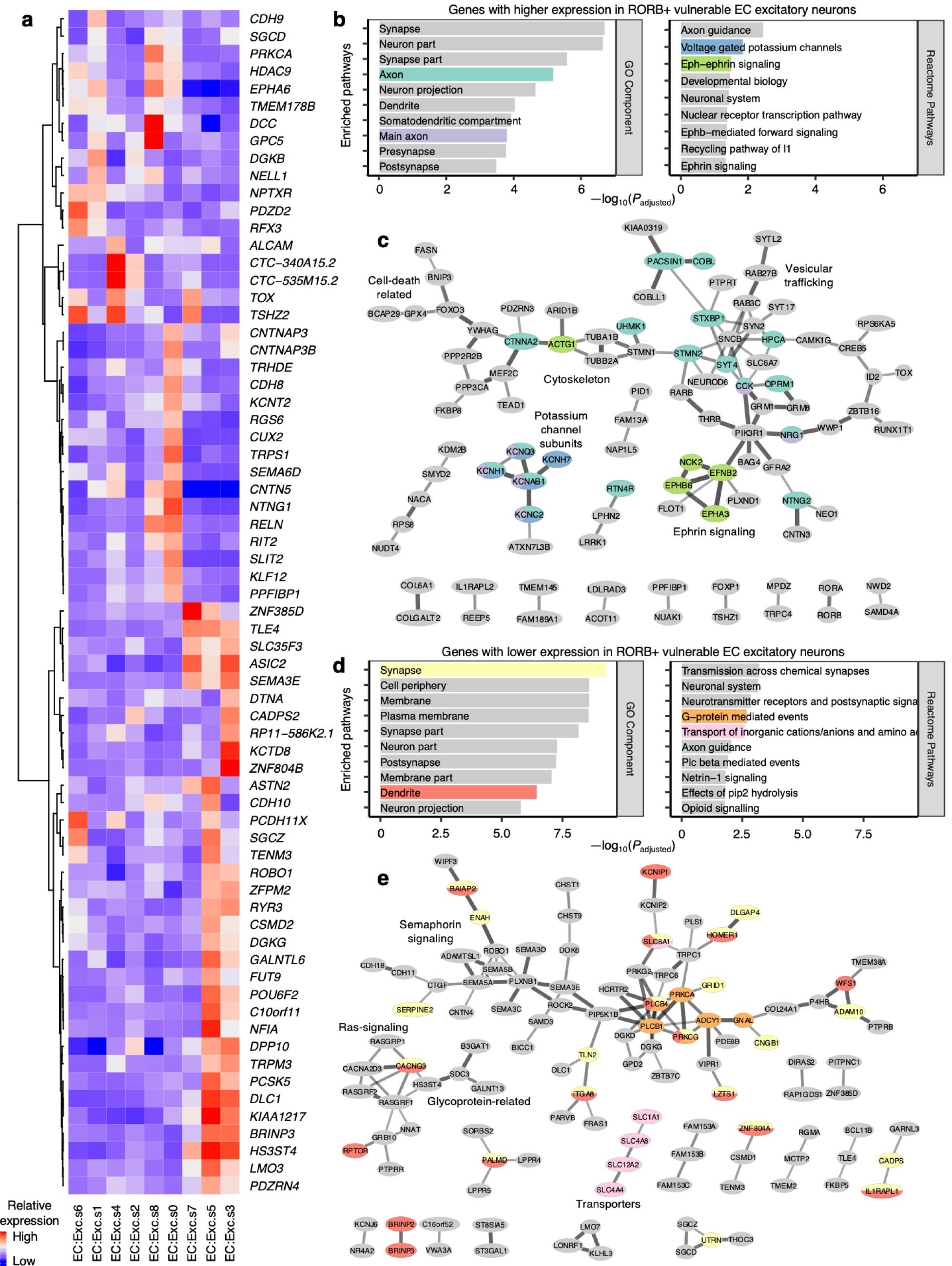
Expression of selected EC excitatory neuron subpopulation markers and pathway enrichment analysis of differentially expressed genes in selectively vulnerable EC excitatory neuron subpopulations. **a**, Expression heatmap of genes that are specifically expressed by four or fewer EC excitatory neuron subpopulations; “High” and “Low” relative expression reflect above- and below-average expression, respectively (see Methods). **b-d**, Enrichment analysis against Gene Ontology Cellular Component terms or Reactome Pathways (**b**,**d**) and functional association network analysis (**c**,**e**; see Methods) of genes with higher (**b-c**) or lower expression (**d-e**) in RORB+ vulnerable EC excitatory neurons, with selected terms highlighted by color. In panels **c** and **e**, genes with stronger associations are connected by thicker lines, and genes without known associations are not shown.

**Extended Data Fig. 3.**
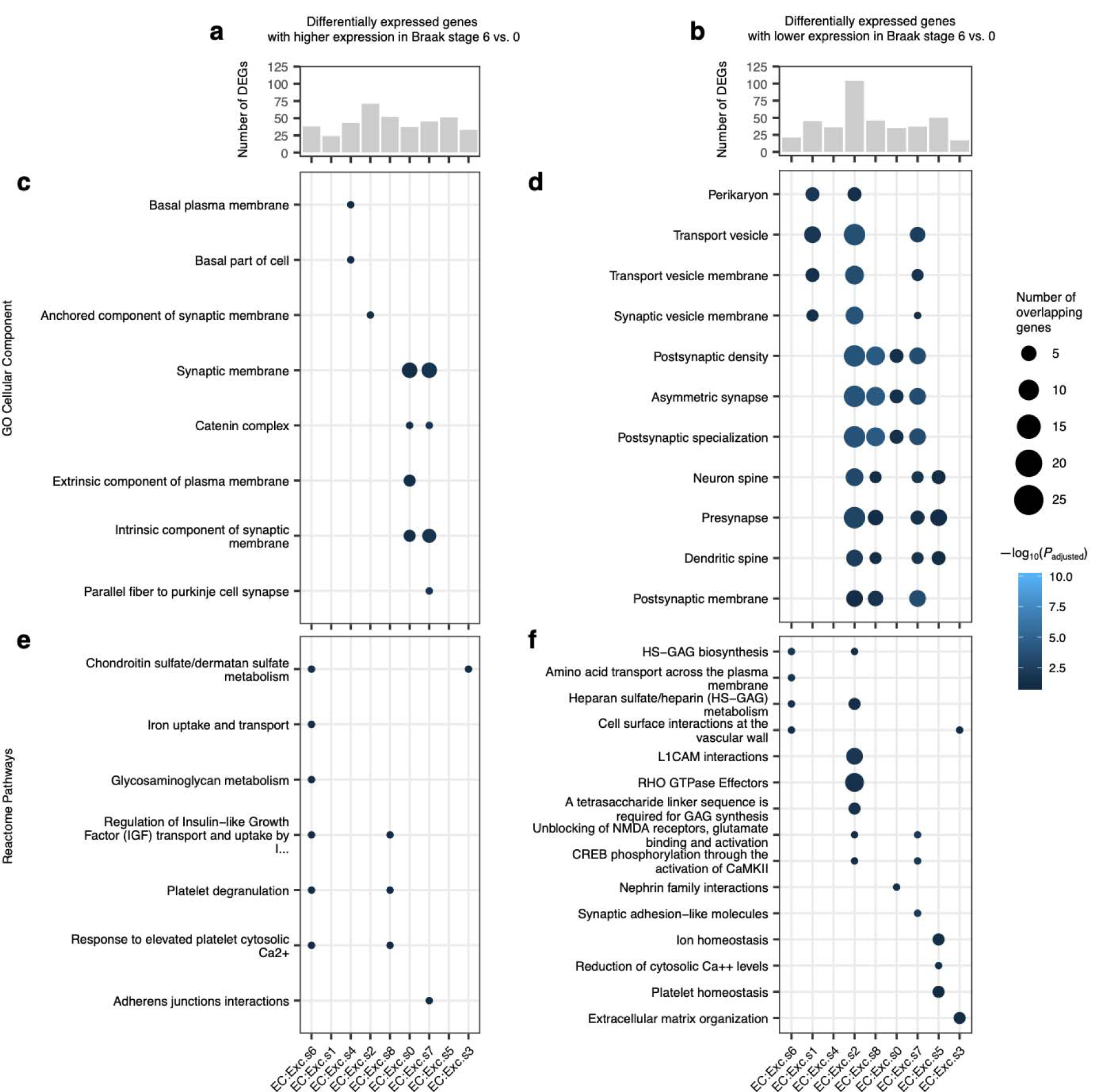
Differential expression analysis across Braak stages for EC excitatory neuron subpopulations. **a-b**, Number of differentially expressed genes in EC excitatory neuron subpopulations with higher (**a**) or lower (**b**) expression in Braak stage 6 vs. Braak stage 0. **c-f**, Enrichment analysis against Gene Ontology Cellular Component terms (**c-d**) or Reactome Pathways (**e-f**) of differentially expressed genes in EC excitatory neuron subpopulations with higher (**c**,**e**) or lower (**d**,**f**) expression in Braak stage 6 vs. Braak stage 0.

**Extended Data Fig. 4.**
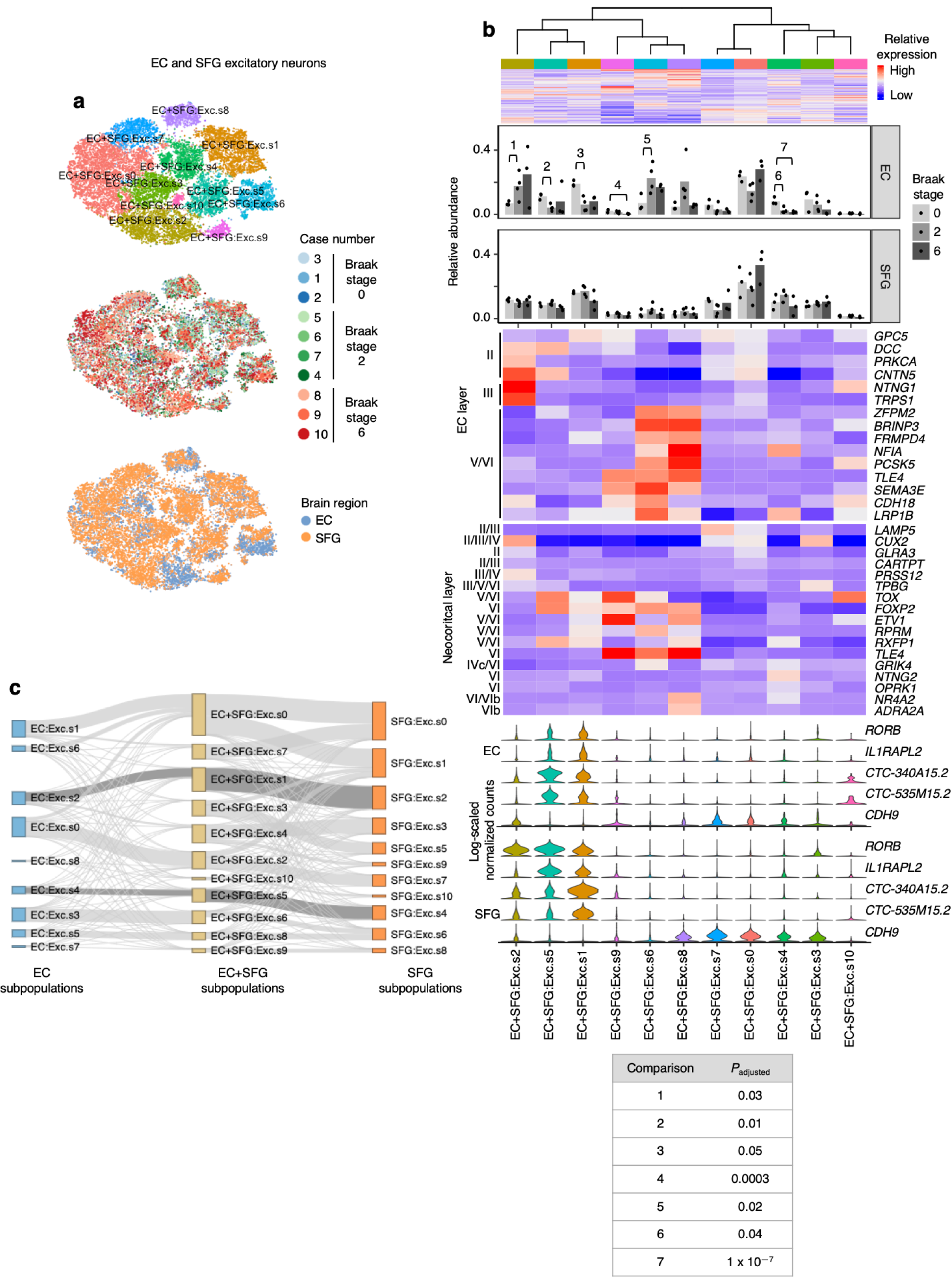
Alignment of EC and SFG maps homologous excitatory neuron subpopulations. **a**, tSNE projection of excitatory neurons from the EC and SFG in the joint alignment space, colored by subpopulation identity (top), individual of origin (middle), or brain region (bottom). **b**, Heatmap and hierarchical clustering of subpopulations and subpopulation marker expression (top subpanel); “High” and “Low” relative expression reflect above- and below-average expression, respectively (see Methods). Relative abundance of subpopulations across Braak stages (second and third subpanels). Expression heatmap of EC layer-specific genes identified from Ramsden *et al*.^34^ (fourth subpanel). Expression heatmap of neocortical layer-specific genes from Lake *et al*.^19^ (fifth subpanel). Expression of selectively vulnerable EC excitatory neuron subpopulation markers by excitatory neurons in the EC (sixth subpanel) or SFG (bottom subpanel). Significant beta regression *P* values (adjusted for multiple testing) are shown in a table at the bottom of the panel. **c**, Sankey diagram connecting subpopulation identity of excitatory neurons in the EC alignment space and the SFG alignment space to subpopulation identity in the EC+SFG alignment space. The links connecting EC:Exc.s2 and EC:Exc.s4 to SFG:Exc.s2 and SFG:Exc.s4, respectively, are highlighted.

**Extended Data Fig. 5.**
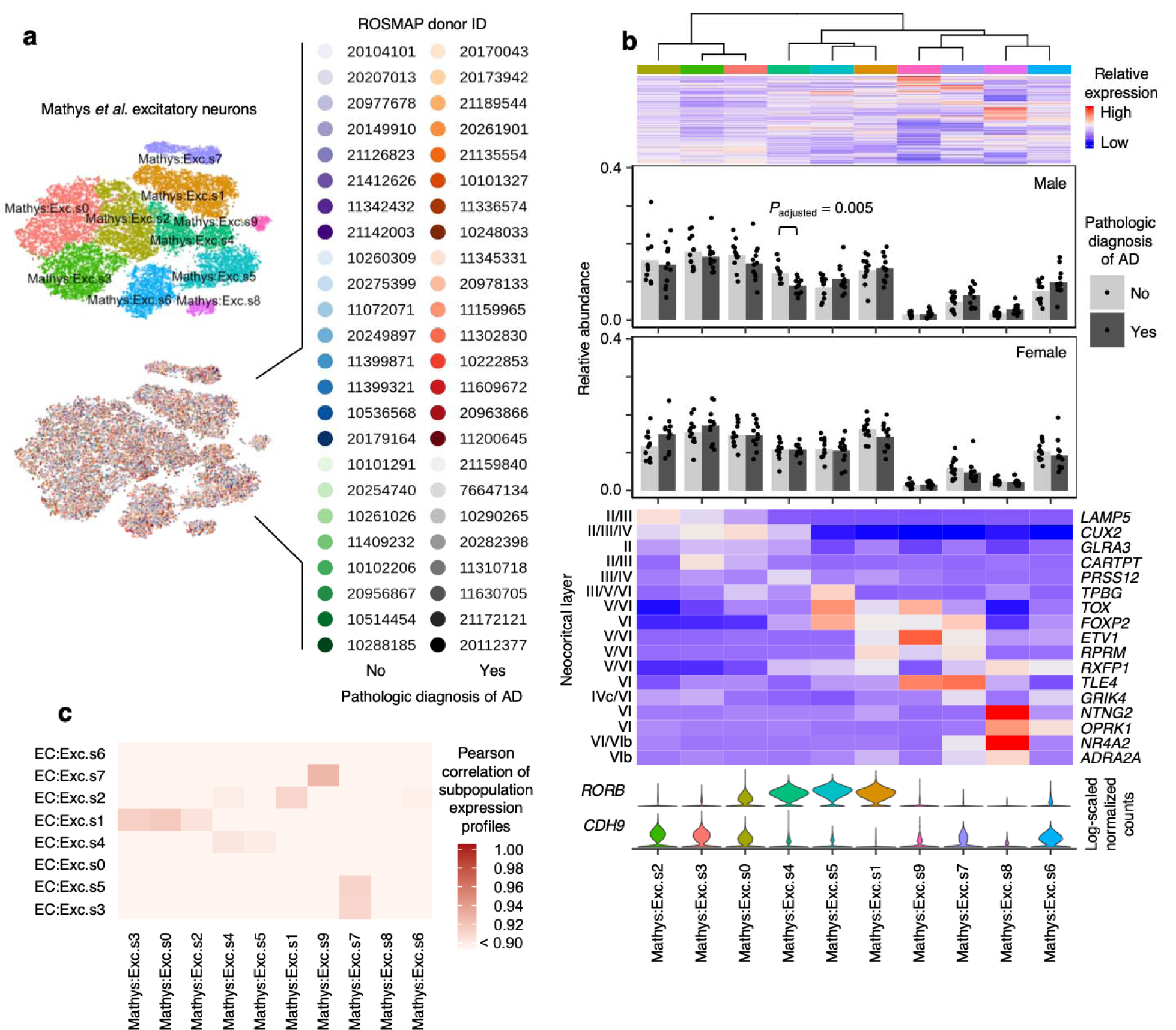
Cross-sample alignment of excitatory neurons from Mathys *et al*. recapitulates selective vulnerability in a RORB-expressing subpopulation. **a**, tSNE projection of excitatory neurons from Mathys *et al*.^21^ in the alignment space, colored by subpopulation identity (top) or individual of origin (bottom). **b**, Heatmap and hierarchical clustering of subpopulations and subpopulation marker expression (top subpanel); “High” and “Low” relative expression reflect above- and below-average expression, respectively (see Methods). Relative abundance of subpopulations in in AD cases vs. controls, separated by sex (second and third subpanels). Expression heatmap of neocortical layer-specific genes from Lake *et al*.^19^ (fourth subpanel). Expression of selectively vulnerable EC excitatory neuron subpopulation markers (bottom subpanel). **c**, Heatmap of Pearson correlation between the gene expression profiles of excitatory neuron subpopulations from the EC vs. those from the prefrontal cortex in Mathys *et al*.^21^

**Extended Data Fig. 6.**
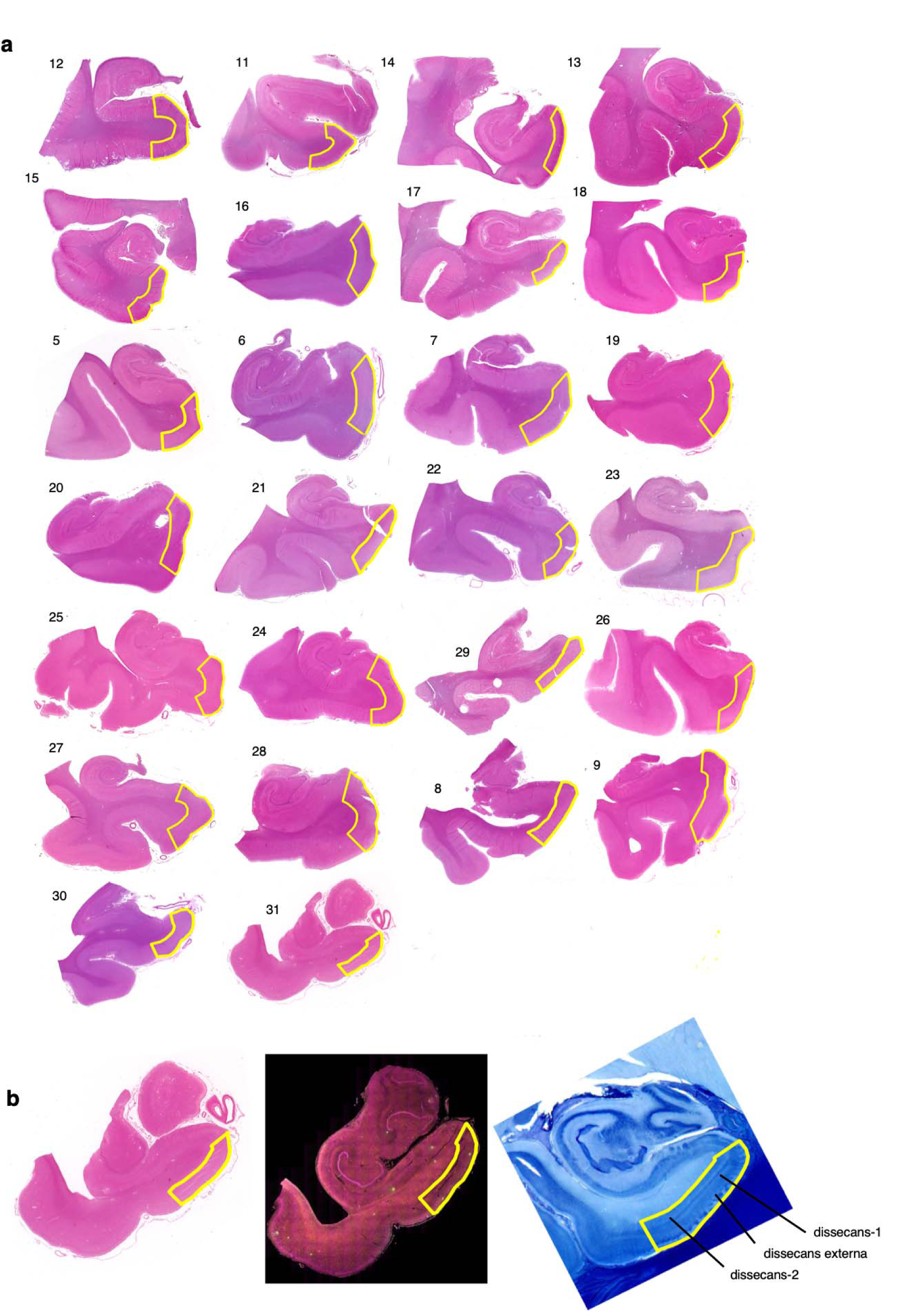
Delineation of the EC for each case used in immunofluorescence validation. **a**, The borders of the caudal EC delineated on sections stained with hematoxylin and eosin (H&E) for all 26 cases used in immunofluorescence validation (Table 1). **b**, Borders of the EC were determined with the aid of 400 um thick serial coronal sections of whole-brain hemispheres stained with gallocyanin (see Methods). Each H&E section (left) along with its corresponding immunofluorescence image (middle) was aligned to the most approximate gallocyanin section (right), in which the the dissecans layers (diss-1, diss-2, and diss-ext) characteristic of the caudal EC were easier to visualize. This was then used to guide delineation of the EC on the H&E and immunofluorescence sections. For more details on the cytoarchitectonic definitions used to define the caudal EC, please consult Heinsen *et al*.^42^.

**Extended Data Fig. 7.**
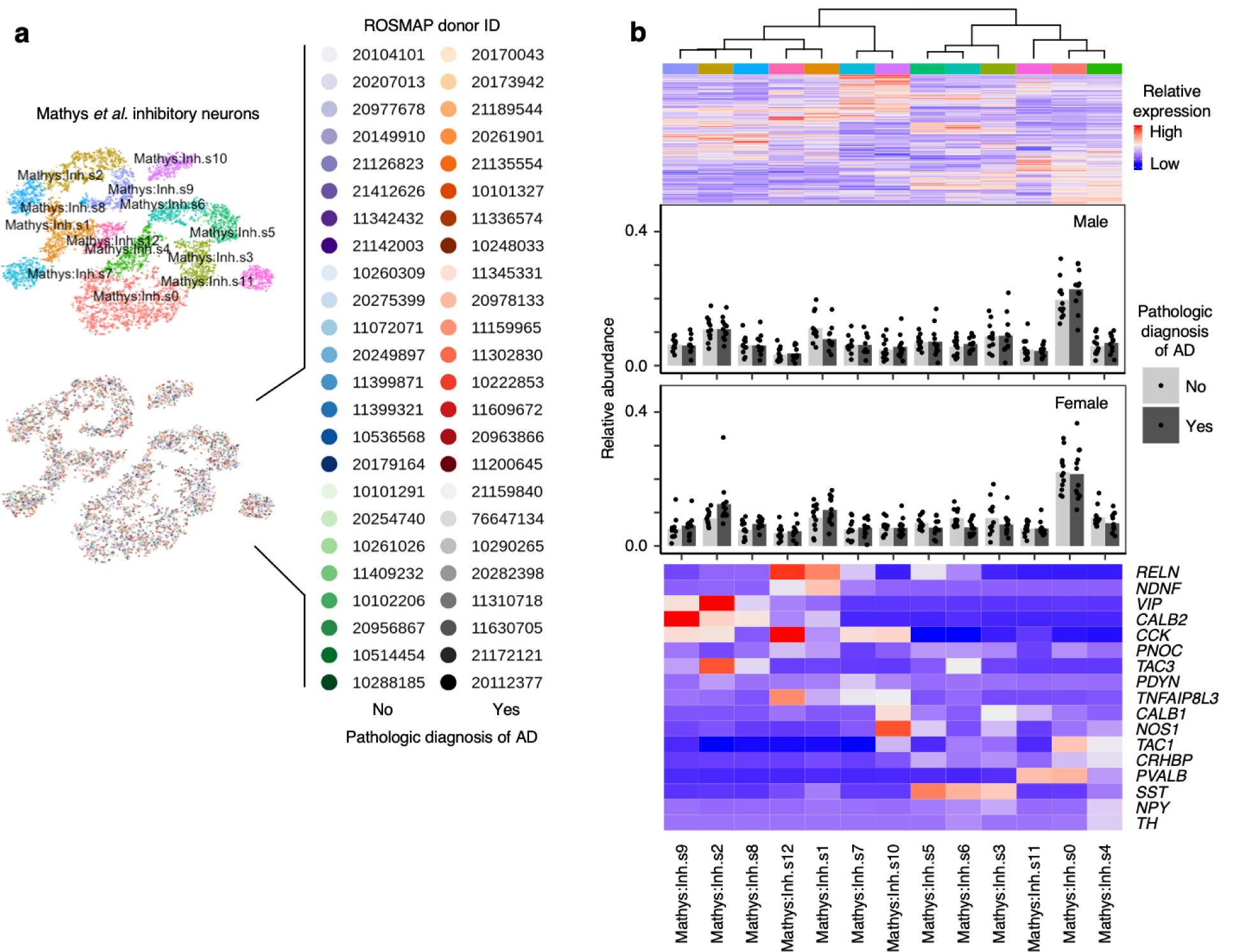
Inhibitory neurons from Mathys *et al*. also do not show differences in resilience or vulnerability to AD. **a**, tSNE projection of inhibitory neurons from Mathys *et al*.^21^ in the alignment space, colored by subpopulation identity (top) or individual of origin (bottom). **b**, Heatmap and hierarchical clustering of subpopulations and subpopulation markers (top subpanel); “High” and “Low” relative expression reflect above- and below-average expression, respectively (see Methods). Relative abundance of subpopulations in in AD cases vs. controls, separated by sex (second and third subpanels). Expression heatmap of inhibitory neuron subtype markers from Lake *et al*.^19^ (bottom subpanel).

**Extended Data Fig. 8.**
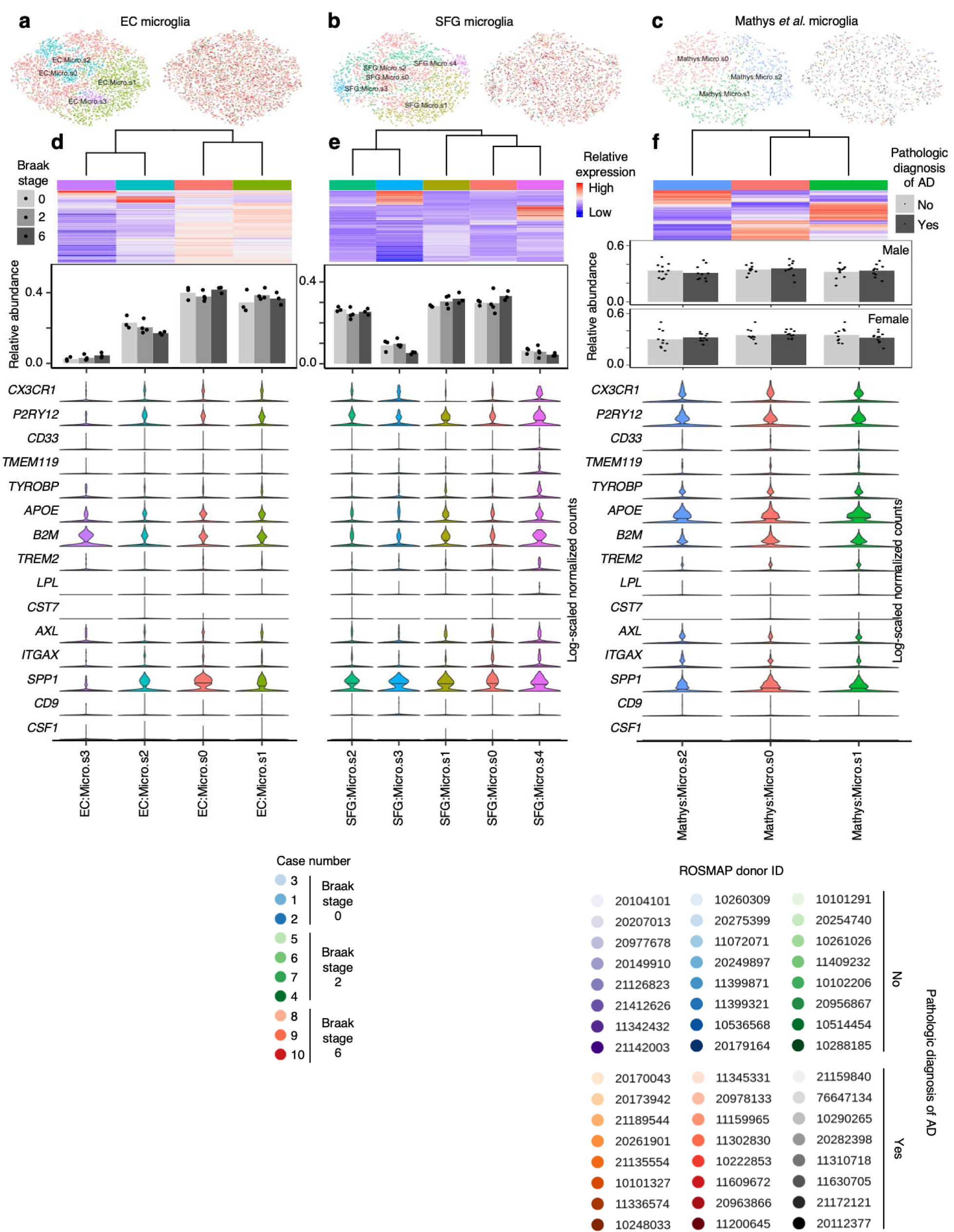
Subclustering of microglia does not sufficiently resolve disease associated microglia signature. **a-c**, tSNE projection of astrocytes from the EC (**a**), SFG (**b**), and Mathys *et al*.^21^ (**c**) in their respective alignment spaces, colored by subpopulation identity (left) or individual of origin (right). **d-f**, Heatmap and hierarchical clustering of subpopulations and subpopulation marker expression (top subpanels); “High” and “Low” relative expression reflect above- and below-average expression, respectively (see Methods). Relative abundance of subpopulations across Braak stages in the EC and SFG or between AD cases vs. controls in Mathys *et al*.^21^ (middle subpanels). Expression of disease associated microglia markers, with median expression level marked by line (bottom subpanels).

**Extended Data Fig. 9.**
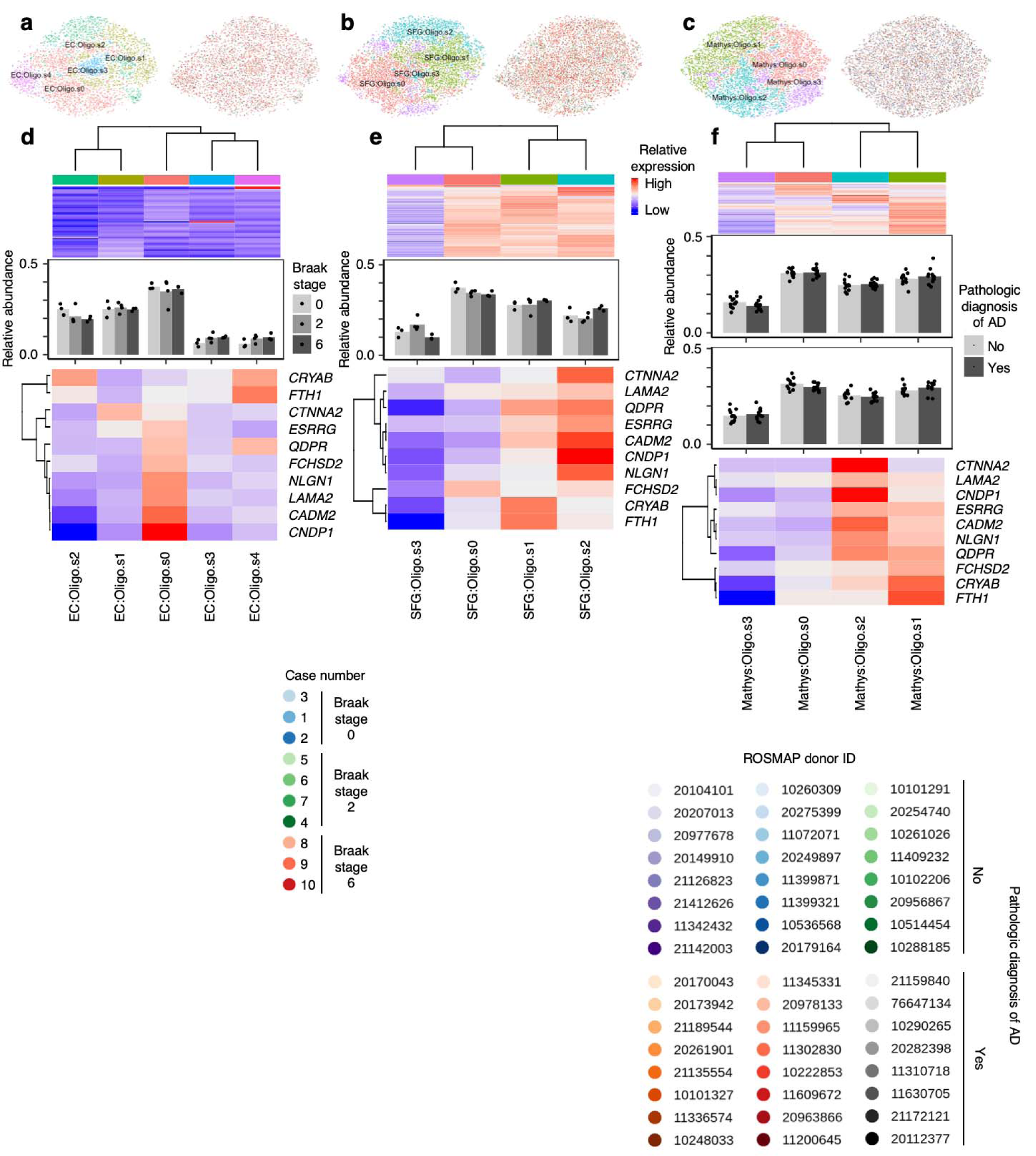
Subclustering of oligodendrocytes identifies subpopulations with higher expression of AD-associated oligodendrocyte markers from Mathys *et al*. **a-c**, tSNE projection of oligodendrocytes from the EC (**a**), SFG (**b**), and Mathys *et al*.^21^ (**c**) in their respective alignment spaces, colored by subpopulation identity (left) or individual of origin (right). **d-f**, Heatmap and hierarchical clustering of subpopulations and subpopulation marker expression (top subpanels); “High” and “Low” relative expression reflect above- and below- average expression, respectively (see Methods). Relative abundance of subpopulations across Braak stages in the EC and SFG or between AD cases vs. controls in Mathys *et al*.^21^ (middle subpanels). Relative expression of AD-associated oligodendrocyte subpopulation markers from Mathys *et al*.^21^ (bottom subpanels).

**Extended Data Fig. 10.**
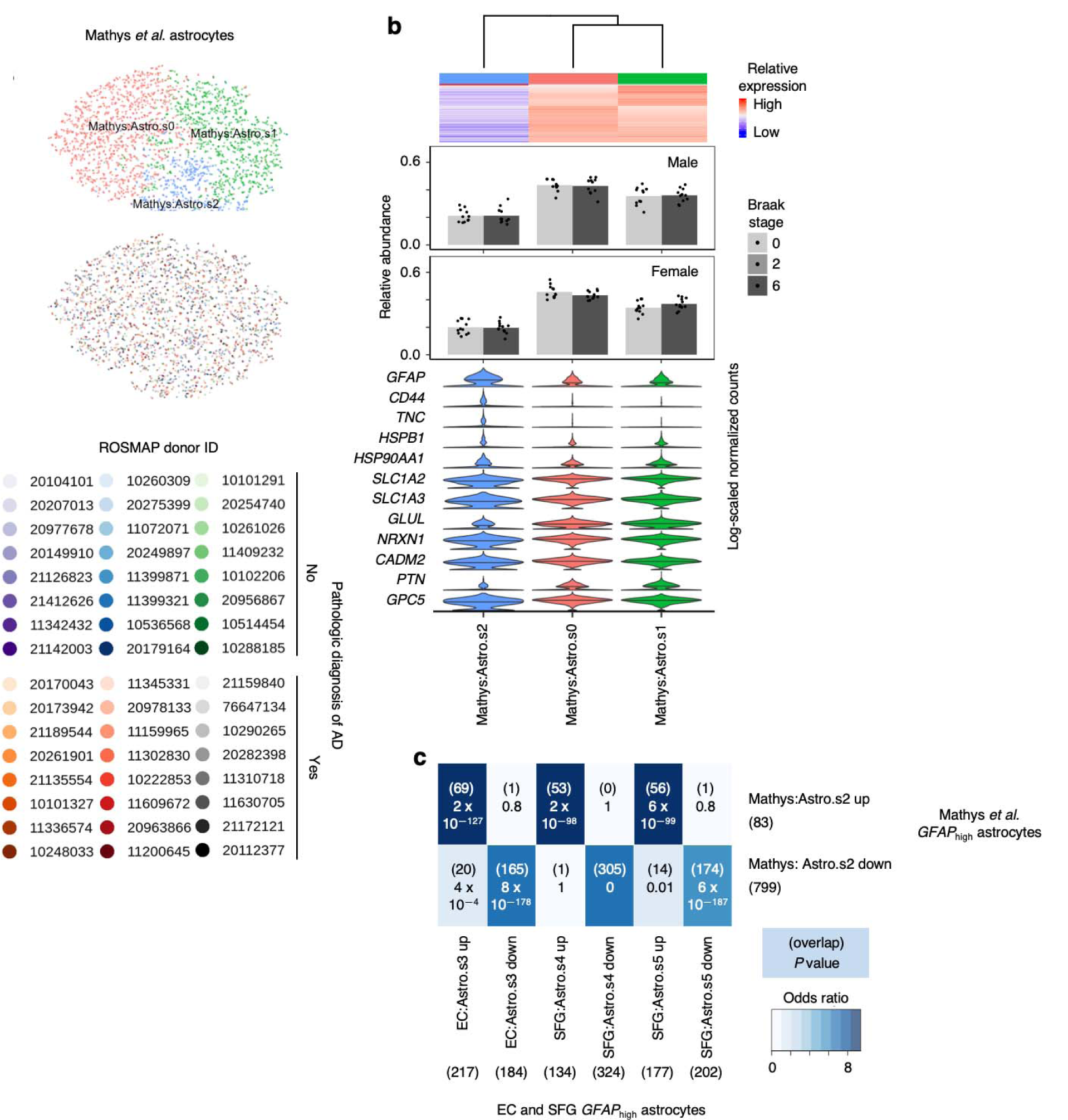
Astrocyte subpopulations with high GFAP expression from Mathys *et al*. are highly similar to those from the EC and SFG. **a**, tSNE projection of astrocytes from Mathys *et al*.^21^ in the alignment subspace, colored by subpopulation identity (top) or individual of origin (bottom). **b**, Heatmap and hierarchical clustering of subpopulations and subpopulation marker expression (top subpanel); “High” and “Low” relative expression reflect above- and below-average expression, respectively (see Methods). Relative abundance of subpopulations in in AD cases vs. controls, separated by sex (middle subpanels). Expression of genes associated with reactive astrocytes, with median expression level marked by line (bottom subpanel). **c**, Enrichment analysis of overlap between differentially expressed genes in astrocytes with high GFAP expression from Mathys *et al*.^21^ vs. differentially expressed genes in astrocytes with high GFAP expression from the EC and SFG; the number of genes in each gene set and the number of overlapping genes are shown in parentheses, and the hypergeometric test p-values are shown without parentheses.

## SUPPLEMENTARY TABLE LEGEND

**Supplemental Table 1** | **Genes differentially expressed by selectively vulnerable excitatory neurons compared to all other excitatory neurons in the EC**. The column “gene” contains official gene symbols of differentially expressed genes, “logFC” contains the associated log_2_ fold-change, “logCPM” contains the log_2_-transformed normalized counts of transcripts mapping to the gene averaged across all conditions, “F” contains the value of the quasi-likelihood F-statistic (see edgeR documentation) used to determine differential expression, “Pvalue” contains the raw *P* values associated with the quasi-likelihood F-test, “FDR” contains *P* values adjusted for multiple testing using the Benjamini-Hochberg method.

## REFERENCES

1. Fu, H., Hardy, J. & Duff, K.E. Selective vulnerability in neurodegenerative diseases. Nat Neurosci 21, 1350–1358 (2018).

2. Saxena, S. & Caroni, P. Selective neuronal vulnerability in neurodegenerative diseases: from stressor thresholds to degeneration. Neuron 71, 35–48 (2011).

3. Braak, H. & Braak, E. Neuropathological stageing of Alzheimer-related changes. Acta Neuropathol 82, 239–259 (1991).

4. Scholl, M., et al. PET Imaging of Tau Deposition in the Aging Human Brain. Neuron 89, 971–982 (2016).

5. Seeley, W.W., Crawford, R.K., Zhou, J., Miller, B.L. & Greicius, M.D. Neurodegenerative diseases target large-scale human brain networks. Neuron 62, 42–52 (2009).

6. Braak, H. & Braak, E. Staging of Alzheimer’s disease-related neurofibrillary changes. Neurobiol Aging 16, 271-278; discussion 278-284 (1995).

7. Price, J.L., et al. Neuron number in the entorhinal cortex and CA1 in preclinical Alzheimer disease. Arch Neurol 58, 1395–1402 (2001).

8. Stranahan, A.M. & Mattson, M.P. Selective vulnerability of neurons in layer II of the entorhinal cortex during aging and Alzheimer’s disease. Neural Plast 2010, 108190 (2010).

9. Van Hoesen, G.W., Hyman, B.T. & Damasio, A.R. Entorhinal cortex pathology in Alzheimer’s disease. Hippocampus 1, 1–8 (1991).

10. Gomez-Isla, T., et al. Profound loss of layer II entorhinal cortex neurons occurs in very mild Alzheimer’s disease. J Neurosci 16, 4491–4500 (1996).

11. Braak, H. & Braak, E. The human entorhinal cortex: normal morphology and lamina-specific pathology in various diseases. Neurosci Res 15, 6–31 (1992).

12. Kordower, J.H., et al. Loss and atrophy of layer II entorhinal cortex neurons in elderly people with mild cognitive impairment. Ann Neurol 49, 202–213 (2001).

13. Gotz, J., Schonrock, N., Vissel, B. & Ittner, L.M. Alzheimer’s disease selective vulnerability and modeling in transgenic mice. Journal of Alzheimer’s disease : JAD 18, 243–251 (2009).

14. Fu, H., et al. Tau Pathology Induces Excitatory Neuron Loss, Grid Cell Dysfunction, and Spatial Memory Deficits Reminiscent of Early Alzheimer’s Disease. Neuron 93, 533–541 e535 (2017).

15. Fu, H., et al. A tau homeostasis signature is linked with the cellular and regional vulnerability of excitatory neurons to tau pathology. Nature neuroscience 22, 47–56 (2019).

16. Drummond, E. & Wisniewski, T. Alzheimer’s disease: experimental models and reality. Acta Neuropathol 133, 155–175 (2017).

17. Liang, W.S., et al. Altered neuronal gene expression in brain regions differentially affected by Alzheimer’s disease: a reference data set. Physiol Genomics 33, 240–256 (2008).

18. Liang, W.S., et al. Gene expression profiles in anatomically and functionally distinct regions of the normal aged human brain. Physiol Genomics 28, 311–322 (2007).

19. Lake, B.B., et al. Neuronal subtypes and diversity revealed by single-nucleus RNA sequencing of the human brain. Science 352, 1586–1590 (2016).

20. Hodge, R.D., et al. Conserved cell types with divergent features in human versus mouse cortex. Nature 573, 61–68 (2019).

21. Mathys, H., et al. Single-cell transcriptomic analysis of Alzheimer’s disease. Nature 570, 332–337 (2019).

22. Grubman, A., et al. A single-cell atlas of entorhinal cortex from individuals with Alzheimer’s disease reveals cell-type-specific gene expression regulation. Nature neuroscience 22, 2087–2097 (2019).

23. Mirra, S.S., et al. The Consortium to Establish a Registry for Alzheimer’s Disease (CERAD). Part II. Standardization of the neuropathologic assessment of Alzheimer’s disease. Neurology 41, 479–486 (1991).

24. Thal, D.R., Rub, U., Orantes, M. & Braak, H. Phases of A beta-deposition in the human brain and its relevance for the development of AD. Neurology 58, 1791–1800 (2002).

25. Braak, H., Thal, D.R., Ghebremedhin, E. & Del Tredici, K. Stages of the pathologic process in Alzheimer disease: age categories from 1 to 100 years. J Neuropathol Exp Neurol 70, 960–969 (2011).

26. Nelson, P.T., et al. Correlation of Alzheimer disease neuropathologic changes with cognitive status: a review of the literature. J Neuropathol Exp Neurol 71, 362–381 (2012).

27. Butler, A., Hoffman, P., Smibert, P., Papalexi, E. & Satija, R. Integrating single-cell transcriptomic data across different conditions, technologies, and species. Nat Biotechnol 36, 411–420 (2018).

28. Haghverdi, L., Lun, A.T.L., Morgan, M.D. & Marioni, J.C. Batch effects in single-cell RNA-sequencing data are corrected by matching mutual nearest neighbors. Nat Biotechnol 36, 421–427 (2018).

29. Johansen, N. & Quon, G. scAlign: a tool for alignment, integration, and rare cell identification from scRNA-seq data. Genome biology 20, 166 (2019).

30. Ferrari, S.L.P. & Cribari-Neto, F. Beta regression for modelling rates and proportions. J Appl Stat 31, 799–815 (2004).

31. Hof, P.R., et al. Parvalbumin-immunoreactive neurons in the neocortex are resistant to degeneration in Alzheimer’s disease. J Neuropathol Exp Neurol 50, 451–462 (1991).

32. Hof, P.R., Cox, K. & Morrison, J.H. Quantitative analysis of a vulnerable subset of pyramidal neurons in Alzheimer’s disease: I. Superior frontal and inferior temporal cortex. J Comp Neurol 301, 44–54 (1990).

33. Hof, P.R. & Morrison, J.H. Neocortical neuronal subpopulations labeled by a monoclonal antabody to calbindin exhibit differential vulnerability in Alzheimer’s disease. Exp Neurol 111, 293–301 (1991).

34. Ramsden, H.L., Surmeli, G., McDonagh, S.G. & Nolan, M.F. Laminar and dorsoventral molecular organization of the medial entorhinal cortex revealed by large-scale anatomical analysis of gene expression. PLoS Comput Biol 11, e1004032 (2015).

35. Kobro-Flatmoen, A. & Witter, M.P. Neuronal chemo-architecture of the entorhinal cortex: A comparative review. Eur J Neurosci 50, 3627–3662 (2019).

36. Naumann, R.K., et al. Conserved size and periodicity of pyramidal patches in layer 2 of medial/caudal entorhinal cortex. J Comp Neurol 524, 783–806 (2016).

37. Jabaudon, D., Shnider, S.J., Tischfield, D.J., Galazo, M.J. & Macklis, J.D. RORbeta induces barrel-like neuronal clusters in the developing neocortex. Cereb Cortex 22, 996–1006 (2012).

38. Oishi, K., Aramaki, M. & Nakajima, K. Mutually repressive interaction between Brn1/2 and Rorb contributes to the establishment of neocortical layer 2/3 and layer 4. Proceedings of the National Academy of Sciences of the United States of America 113, 3371–3376 (2016).

39. Nakagawa, Y. & O’Leary, D.D. Dynamic patterned expression of orphan nuclear receptor genes RORalpha and RORbeta in developing mouse forebrain. Dev Neurosci 25, 234–244 (2003).

40. Marinaro, F., et al. Molecular and cellular pathology of monogenic Alzheimer’s disease at single cell resolution. bioRxiv, 2020.2007.2014.202317 (2020).

41. Franjic, D., et al. Molecular Diversity Among Adult Human Hippocampal and Entorhinal Cells. bioRxiv doi: https://doi.org/10.1101/2019.12.31.889139 (2019).

42. Heinsen, H., et al. Quantitative investigations on the human entorhinal area: left-right asymmetry and age-related changes. Anat Embryol (Berl) 190, 181–194 (1994).

43. Ehrenberg, A.J., et al. A manual multiplex immunofluorescence method for investigating neurodegenerative diseases. Journal of Neuroscience Methods In press, https://doi.org/10.1016/j.jneumeth.2020.108708. (2020).

44. Keren-Shaul, H., et al. A Unique Microglia Type Associated with Restricting Development of Alzheimer’s Disease. Cell 169, 1276–1290 e1217 (2017).

45. Srinivasan, K., et al. Alzheimer’s patient brain myeloid cells exhibit enhanced aging and unique transcriptional activation. bioRxiv doi: https://doi.org/10.1101/610345 (2019).

46. Thrupp, N., et al. Single nucleus sequencing fails to detect microglial activation in human tissue. bioRxiv, 2020.2004.2013.035386 (2020).

47. Chen, W.T., et al. Spatial Transcriptomics and In Situ Sequencing to Study Alzheimer’s Disease. Cell (2020).

48. Chun, H. & Lee, C.J. Reactive astrocytes in Alzheimer’s disease: A double-edged sword. Neurosci Res 126, 44–52 (2018).

49. Perez-Nievas, B.G. & Serrano-Pozo, A. Deciphering the Astrocyte Reaction in Alzheimer’s Disease. Front Aging Neurosci 10, 114 (2018).

50. Simpson, J.E., et al. Microarray analysis of the astrocyte transcriptome in the aging brain: relationship to Alzheimer’s pathology and APOE genotype. Neurobiol Aging 32, 1795–1807 (2011).

51. Sekar, S., et al. Alzheimer’s disease is associated with altered expression of genes involved in immune response and mitochondrial processes in astrocytes. Neurobiol Aging 36, 583–591 (2015).

52. Liddelow, S.A., et al. Neurotoxic reactive astrocytes are induced by activated microglia. Nature 541, 481–487 (2017).

53. Brodkey, J.A., et al. Focal brain injury and upregulation of a developmentally regulated extracellular matrix protein. J Neurosurg 82, 106–112 (1995).

54. Laywell, E.D., et al. Enhanced expression of the developmentally regulated extracellular matrix molecule tenascin following adult brain injury. Proceedings of the National Academy of Sciences of the United States of America 89, 2634–2638 (1992).

55. Zamanian, J.L., et al. Genomic analysis of reactive astrogliosis. J Neurosci 32, 6391–6410 (2012).

56. Anderson, M.A., et al. Astrocyte scar formation aids central nervous system axon regeneration. Nature 532, 195–200 (2016).

57. Insausti, R., Munoz-Lopez, M., Insausti, A.M. & Artacho-Perula, E. The Human Periallocortex: Layer Pattern in Presubiculum, Parasubiculum and Entorhinal Cortex. A Review. Front Neuroanat 11, 84 (2017).

58. Mikkonen, M., Alafuzoff, I., Tapiola, T., Soininen, H. & Miettinen, R. Subfield- and layer-specific changes in parvalbumin, calretinin and calbindin-D28K immunoreactivity in the entorhinal cortex in Alzheimer’s disease. Neuroscience 92, 515–532 (1999).

59. Winterer, J., et al. Excitatory Microcircuits within Superficial Layers of the Medial Entorhinal Cortex. Cell Rep 19, 1110–1116 (2017).

60. Braak, H. & Braak, E. On areas of transition between entorhinal allocortex and temporal isocortex in the human brain. Normal morphology and lamina-specific pathology in Alzheimer’s disease. Acta Neuropathol 68, 325–332 (1985).

61. Kampmann, M. A CRISPR Approach to Neurodegenerative Diseases. Trends in molecular medicine 23, 483–485 (2017).

62. Kampmann, M. CRISPR-based functional genomics for neurological disease. Nat Rev Neurol (2020).

63. Tian, R., et al. CRISPR Interference-Based Platform for Multimodal Genetic Screens in Human iPSC-Derived Neurons. Neuron 104, 239–255 e212 (2019).

64. Tian, R., et al. CRISPR Interference-Based Platform for Multimodal Genetic Screens in Human iPSC-Derived Neurons. Neuron 104, 239–255 (2019).

65. Grinberg, L.T., et al. Brain bank of the Brazilian aging brain study group - a milestone reached and more than 1,600 collected brains. Cell Tissue Bank 8, 151–162 (2007).

66. Hyman, B.T., et al. National Institute on Aging-Alzheimer’s Association guidelines for the neuropathologic assessment of Alzheimer’s disease. Alzheimers Dement 8, 1–13 (2012).

67. Suemoto, C.K., et al. Neuropathological diagnoses and clinical correlates in older adults in Brazil: A cross-sectional study. PLoS Med 14, e1002267 (2017).

68. Cairns, N.J., et al. Neuropathologic diagnostic and nosologic criteria for frontotemporal lobar degeneration: consensus of the Consortium for Frontotemporal Lobar Degeneration. Acta Neuropathol 114, 5–22 (2007).

69. Ferrer, I., Santpere, G. & van Leeuwen, F.W. Argyrophilic grain disease. Brain 131, 1416–1432 (2008).

70. Rodriguez, R.D. & Grinberg, L.T. Argyrophilic grain disease: An underestimated tauopathy. Dement Neuropsychol 9, 2–8 (2015).

71. Rodriguez, R.D., et al. Argyrophilic Grain Disease: Demographics, Clinical, and Neuropathological Features From a Large Autopsy Study. J Neuropathol Exp Neurol 75, 628–635 (2016).

72. Mo, A., et al. Epigenomic Signatures of Neuronal Diversity in the Mammalian Brain. Neuron 86, 1369–1384 (2015).

73. Tabula Muris, C., et al. Single-cell transcriptomics of 20 mouse organs creates a Tabula Muris. Nature 562, 367–372 (2018).

74. Lun, A.T.L., et al. EmptyDrops: distinguishing cells from empty droplets in droplet-based single-cell RNA sequencing data. Genome biology 20, 63 (2019).

75. Lun, A.T., Bach, K. & Marioni, J.C. Pooling across cells to normalize single-cell RNA sequencing data with many zero counts. Genome biology 17, 75 (2016).

76. Lun, A.T., McCarthy, D.J. & Marioni, J.C. A step-by-step workflow for low-level analysis of single-cell RNA-seq data with Bioconductor. F1000Res 5, 2122 (2016).

77. McCarthy, D.J., Campbell, K.R., Lun, A.T. & Wills, Q.F. Scater: pre-processing, quality control, normalization and visualization of single-cell RNA-seq data in R. Bioinformatics 33, 1179–1186 (2017).

78. Stuart, T., et al. Comprehensive Integration of Single-Cell Data. Cell 177, 1888–1902 e1821 (2019).

79. Robinson, M.D., McCarthy, D.J. & Smyth, G.K. edgeR: a Bioconductor package for differential expression analysis of digital gene expression data. Bioinformatics 26, 139–140 (2010).

80. Amezquita, R.A., et al. Orchestrating single-cell analysis with Bioconductor. Nat Methods 17, 137–145 (2020).

81. Szklarczyk, D., et al. STRING v11: protein-protein association networks with increased coverage, supporting functional discovery in genome-wide experimental datasets. Nucleic acids research 47, D607–D613 (2019).

82. Shannon, P., et al. Cytoscape: a software environment for integrated models of biomolecular interaction networks. Genome Res 13, 2498–2504 (2003).

83. Cribari-Neto, F. & Zeileis, A. Beta Regression in R. Journal of Statistical Software 34, 1–24 (2010).

84. Stelzer, G., et al. The GeneCards Suite: From Gene Data Mining to Disease Genome Sequence Analyses. Curr Protoc Bioinformatics 54, 1 30 31–31 30 33 (2016).

85. Arriza, J.L., et al. Functional comparisons of three glutamate transporter subtypes cloned from human motor cortex. J Neurosci 14, 5559–5569 (1994).

86. Borden, L.A., et al. Cloning of the human homologue of the GABA transporter GAT-3 and identification of a novel inhibitor with selectivity for this site. Receptors Channels 2, 207–213 (1994).

87. Gendreau, S., et al. A trimeric quaternary structure is conserved in bacterial and human glutamate transporters. J Biol Chem 279, 39505–39512 (2004).

88. Häberle, J., et al. Congenital glutamine deficiency with glutamine synthetase mutations. N Engl J Med 353, 1926–1933 (2005).

89. Kawakami, H., Tanaka, K., Nakayama, T., Inoue, K. & Nakamura, S. Cloning and expression of a human glutamate transporter. Biochem Biophys Res Commun 199, 171–176 (1994).

90. Melzer, N., Biela, A. & Fahlke, C. Glutamate modifies ion conduction and voltage-dependent gating of excitatory amino acid transporter-associated anion channels. J Biol Chem 278, 50112–50119 (2003).

91. Südhof, T.C. Synaptic Neurexin Complexes: A Molecular Code for the Logic of Neural Circuits. Cell 171, 745–769 (2017).

92. Pellissier, F., Gerber, A., Bauer, C., Ballivet, M. & Ossipow, V. The adhesion molecule Necl-3/SynCAM-2 localizes to myelinated axons, binds to oligodendrocytes and promotes cell adhesion. BMC Neurosci 8, 90 (2007).

93. González-Castillo, C., Ortuño-Sahagún, D., Guzmán-Brambila, C., Pallàs, M. & Rojas-Mayorquín, A.E. Pleiotrophin as a central nervous system neuromodulator, evidences from the hippocampus. Front Cell Neurosci 8, 443 (2014).

94. Siddiqui, T.J., et al. An LRRTM4-HSPG complex mediates excitatory synapse development on dentate gyrus granule cells. Neuron 79, 680–695 (2013).

95. Heinsen, H., Arzberger, T. & Schmitz, C. Celloidin mounting (embedding without infiltration) - a new, simple and reliable method for producing serial sections of high thickness through complete human brains and its application to stereological and immunohistochemical investigations. J Chem Neuroanat 20, 49–59 (2000).

96. Insausti, R. & Amaral, D.G. Entorhinal cortex of the monkey: IV. Topographical and laminar organization of cortical afferents. J Comp Neurol 509, 608–641 (2008).

97. Rose, S. Vergleichende Messungen im Allocortex bei Tier und Mensch. J Psychol Neurol 34, 250–255 (1927).

98. Hevner, R.F., et al. Tbr1 regulates differentiation of the preplate and layer 6. Neuron 29, 353–366 (2001).

99. Ehrenberg, A.J., et al. Quantifying the accretion of hyperphosphorylated tau in the locus coeruleus and dorsal raphe nucleus: the pathological building blocks of early Alzheimer’s disease. Neuropathology and applied neurobiology 43, 393–408 (2017).

100. Montine, T.J., et al. National Institute on Aging-Alzheimer’s Association guidelines for the neuropathologic assessment of Alzheimer’s disease: a practical approach. Acta Neuropathol 123, 1–11 (2012).

101. Hughes, C.P., Berg, L., Danziger, W.L., Coben, L.A. & Martin, R.L. A new clinical scale for the staging of dementia. Br J Psychiatry 140, 566–572 (1982).

